# Pore dynamics and asymmetric cargo loading in an encapsulin nanocompartment

**DOI:** 10.1101/2021.04.15.439977

**Authors:** Jennifer Ross, Zak McIver, Thomas Lambert, Cecilia Piergentili, Jasmine Emma Bird, Kelly J. Gallagher, Faye L. Cruickshank, Patrick James, Efrain Zarazúa-Arvizu, Louise E. Horsfall, Kevin J. Waldron, Marcus D. Wilson, C. Logan Mackay, Arnaud Baslé, David J. Clarke, Jon Marles-Wright

## Abstract

Encapsulins are protein nanocompartments that house various cargo enzymes, including a family of decameric ferritin-like proteins. Here, we study a recombinant *Haliangium ochraceum* encapsulin:encapsulated ferritin complex using electron cryo-microscopy and hydrogen/deuterium exchange mass spectrometry to gain insight into the structural relationship between the encapsulin shell and its protein cargo. An asymmetric single particle reconstruction reveals four encapsulated ferritin decamers in a tetrahedral arrangement within the encapsulin nanocompartment. This leads to a symmetry mismatch between the protein cargo and the icosahedral encapsulin shell. The encapsulated ferritin decamers are offset from the interior face of the encapsulin shell. Using HDX-MS, we observed dynamic behavior of the major five-fold pore in the encapsulin shell and show the pore opening via the movement of the encapsulin A-domain. These data will accelerate efforts to engineer the encapsulation of heterologous cargo proteins and to alter the permeability of the encapsulin shell via pore modifications.

**Teaser:** Cryo-EM and HDX-MS analysis of an encapsulin nanocompartment shows that the pores at the five-fold icosahedral vertex of the shell are flexible.

## Introduction

Cellular metabolism and reaction pathways can produce toxic by-products which damage proteins, DNA, and lipids, or can become involved in potentially harmful side-reactions. Eukaryotes use membrane-bound organelles, such as lysosomes, to prevent this damage by housing dangerous reactions in chemically privileged environments. In a similar manner, prokaryotes use large protein-based compartments to sequester such reactions and act as barrier from the cytosol (*1, 2*). Prokaryotes have a variety of metabolic compartments including carboxysomes, which are used for carbon dioxide fixation in photosynthetic and some chemotrophic bacteria(*3, 4*); the related bacterial microcompartments, which allow the catabolism of carbon sources that produce aldehyde intermediates in their breakdown, such as ethanolamine and propanediol(*5*); and ferritins, used for iron oxidation and storage (*6, 7*).

One other key compartmentalization strategy utilized by prokaryotes and archaea is the encapsulin system (*8*–*10*). Encapsulin (Enc) nanocompartments are hollow icosahedral complexes which range in size from 20 nm to 42 nm (*9, 11, 12*). Encapsulin proteins are structurally related to the viral capsid protein (gp5) of the HK97 bacteriophage and self-assemble from a single monomer into one of three forms: 60 subunits (*T* = 1 capsid symmetry), 180 subunits (*T* = 3 capsid symmetry) or 240 subunits (*T* = 4 symmetry)(*8, 9, 12, 13*). Encapsulins share a common feature of housing a cargo enzyme, such as ferritin-like proteins (encapsulated ferritins, EncFtn), iron-mineralizing encapsulin-associated firmicute (IMEF), or dye-decolorizing peroxidases(*12, 14*). Cargo enzymes are directed inside the encapsulin nanocompartment by a terminal localization sequence (LS) which binds to the interior face of the encapsulin(*9, 15*). Encapsulins and their cargo proteins are found throughout the bacterial and archaeal domains in species inhabiting a range of environmental niches; consequently, the proteins are stable in diverse physical conditions(*8, 16*–*19*). For these reasons, the encapsulins have attracted considerable interest for biotechnological applications, through their ability to separate potentially hazardous heterologous reactions from the native cytosol(*20, 21*). The expression of functional engineered encapsulins has also been demonstrated in eukaryotes, where they have been developed as metabolic organelles in *Saccharomyces cerevisiae*(*21*) and in cellular imaging applications in human cells(*22*).

The EncFtn cargo proteins are of particular interest, as they differ from their classical ferritin relatives. Although both proteins oxidize iron using a conserved catalytically active ferroxidase center (FOC), they have remarkably different structural architectures. Classical ferritins oxidize ferrous iron, Fe(II), into a mineral ferric form, Fe(III), which is then stored within a 24-meric 12 nm nanocage (*6, 23*). In contrast to this, the EncFtn proteins have an annular structure formed from a pentamer of dimers with the FOC active sites located at a dimer interface (*11, 16*). EncFtn oxidizes iron in a similar manner to other ferritins, but due to its open structure, it must be associated with an encapsulin nanocage to act as an iron store (*11*). Together the encapsulin EncFtn (Enc:EncFtn) complex can perform both the oxidation and storage functions of classical ferritins. However, due to its increased size when compared to classical ferritins, the Enc:EncFtn complex has the potential to house significantly greater quantities of iron, and has been described as an iron megastore (*13*). For these reasons, the encapsulins have attracted considerable interest for biotechnological applications, through their ability to separate potentially hazardous heterologous reactions from the host cytosol.

Although there have been several structural studies on encapsulins, several key questions remain unanswered. Most notably, for the EncFtn containing encapsulin nanocompartments, the structural relationship between the encapsulin shell and the EncFtn cargo protein is unknown. Studies on Dyp loaded encapsulin nanocompartments have shown loading with either one(*24*) or two hexameric complexes(*25*). Previous models for the loading of EncFtn in encapsulin nanocompartments have suggested a symmetric arrangement of the D5 decameric encapsulated ferritin at the five-fold icosahedral vertices of the encapsulin shell, giving a theoretical maximum of twelve EncFtn decamers per encapsulin nanocage(*11, 26*).

Herein, we investigate the structure of the Enc:EncFtn nanocompartment from the halophilic bacterium *Haliangium ochraceum*, to gain a better understanding of the arrangement and stoichiometry of the complex. Using the complementary structural biology techniques of cryogenic electron microscopy (cryo-EM) and hydrogen-deuterium exchange mass spectrometry (HDX-MS), we study the structural relationship between Enc and EncFtn in the encapsulin nanocage.

We present the first cryo-EM structure of a mesophilic Enc:EncFtn nanocompartment, which reveals a symmetry-breaking tetrahedral arrangement of the EncFtn decamers within the encapsulin shell. Analysis of the encapsulin shell by symmetry expansion of an icosahedral reconstructions and focused 3D refinement on the pentameric vertex reveals a flexible pore in the shell. The dynamic nature of this pore region was investigated by HDX-MS. We show that this region has a high rate of H/D exchange, demonstrating its conformational flexibility. Our combination of HDX-MS and cryo-EM models affords insight into the loading capacity and dynamics of the Enc:EncFtn nanocompartment system.

## Results

### Recombinant *Haliangium ochraceum* encapsulin complexes form regular nanocompartments that recruit active EncFtn cargoes

In order to gain an understanding of the relationship between encapsulin nanocompartments and their EncFtn cargoes, the Enc:EncFtn nanocompartment from the halophilic mesophile *Haliangium ochraceum* was chosen as our model system. This was primarily due to the high yields and ease of purification of the recombinant nanocompartment. Constructs to produce empty (Empty-Enc) and EncFtn loaded (Loaded-Enc) encapsulin nanocompartments were assembled for recombinant protein expression in *Escherichia coli*. The protein complexes were purified by heat treatment, followed by anion exchange and size-exclusion chromatography (**Figure 1A and B, and Figure S1A-D**).

**Figure 1.**
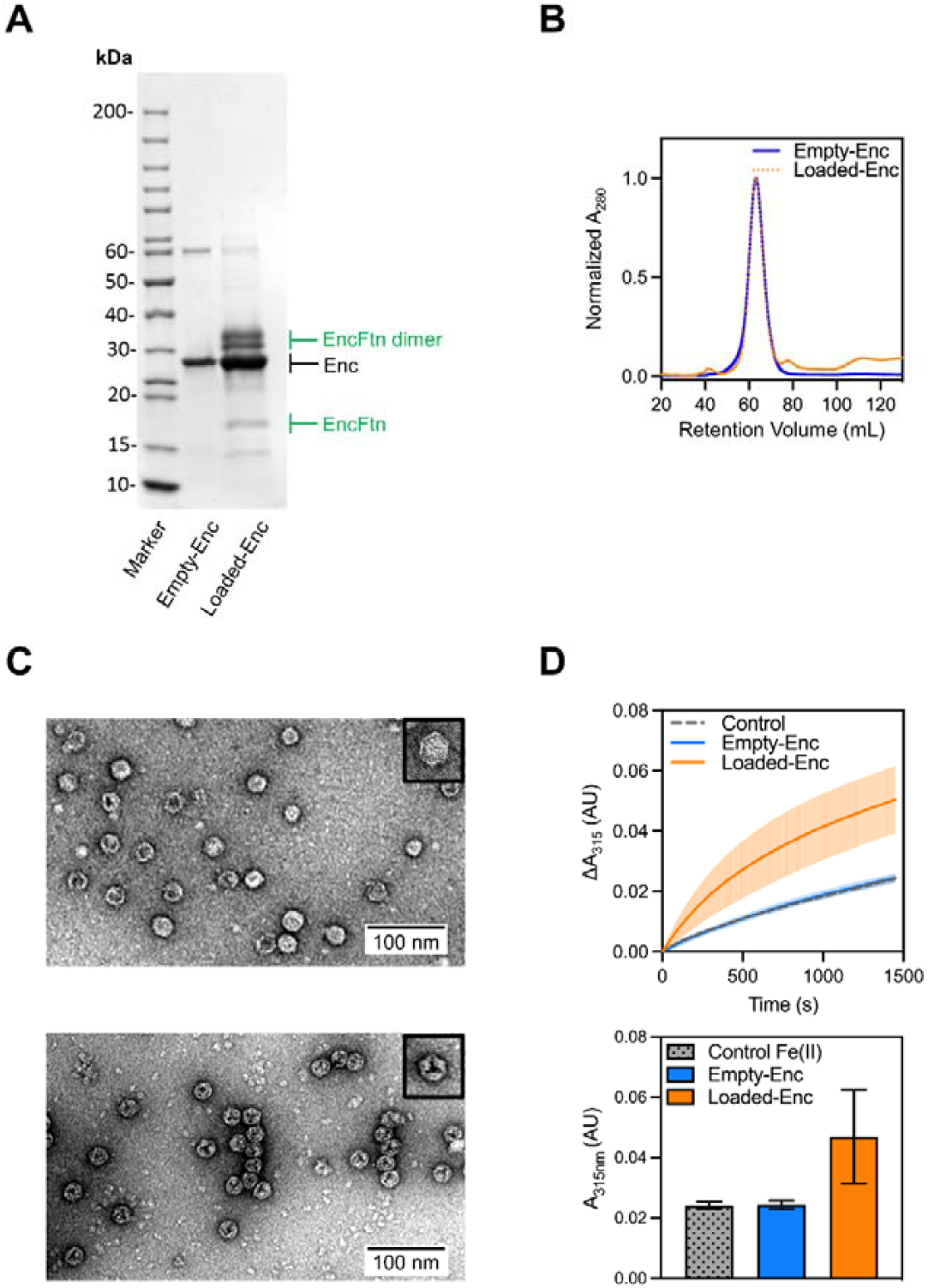
Validation of the Assembly and Activity of Loaded-Enc and Empty-Enc nanocompartments. **A**: SDS-PAGE of purified Empty-Enc and Loaded-Enc. Proteins resolved by 15% acrylamide SDS-PAGE and stained with Coomassie blue stain. Encapsulin bands are near the 30 kDa marker and highlighted by a black arrow. The EncFtn cargo of Loaded-Enc appears as both a monomer and dimer. Bands corresponding to the monomer and dimer of EncFtn are highlighted with green arrows. **B**: Recombinant, purified Empty-Enc (blue trace) and Loaded-Enc (orange, dotted trace) were analyzed by size exclusion chromatography using a Sephacryl 400 column (Cytiva). Both encapsulins elute at the same volume, suggesting the encapsulin complexes are the same size regardless of cargo loading. **C**: Negative stain transmission micrographs of Empty-Enc (upper micrograph) and Loaded-Enc (lower micrograph) displaying individual particles for each complex. One nanocompartment of Empty-Enc and Loaded-Enc is shown in the upper right corner of each micrograph, with a hexagonal 2D geometry observed. **D**: upper panel: Ferroxidase activity of Loaded-Enc compared to Empty-Enc. Protein samples were mixed with 100 μM FeSO_4_.7H_2_O. Following an incubation period at room temperature of 50 seconds absorbance at 315 nm was measured over a time-course of 1450 seconds. Control reference established using enzyme-free reaction as a measure of background iron oxidation. The lines represent the mean of all repeats, error bars represent the standard deviation from the mean. Lower panel: End point ferroxidase assay comparison. Ferroxidase activity shown by the total increase in A_315 nm_ at the end point of the assay. Bars represent the mean of all repeats, error bars represent the standard deviation from the mean.

Size-exclusion chromatography was used to separate correctly formed encapsulin nanocompartments from contaminating proteins and partially formed sub-complexes (**Figure S1A/B**). Fractions corresponding to the major peaks were visualized by negative stain transmission electron microscopy (TEM) (**Figure S2**), with the fraction corresponding to a retention volume of 63 mL for both the Empty-Enc and Loaded-Enc containing the homogeneous encapsulin nanocompartments (**Figure 1C**). Both empty and EncFtn-loaded encapsulins assembled into regular nanocompartments, with an average diameter of approximately 21 nm, consistent with other *T=*1 type encapsulins (**Figure S1E and F**) (*9, 25, 27, 28*). The micrographs of the Loaded-Enc sample reveal a regular internal density visible within the nanocompartment, suggesting that the EncFtn cargo has been encapsulated in an organized manner.

The molecular masses of the protein constituents of the Empty-Enc and Loaded-Enc nanocompartments were determined by LC-MS (**Table S1**). MS analysis of the Empty-Enc assembly revealed a single charge state distribution corresponding to a monomer of the encapsulin protein. MS analysis of the Loaded-Enc revealed three charge state distributions present, with deconvoluted masses consistent with the encapsulin protein monomer, a monomer of EncFtn and a dimer of EncFtn (**Figure S1G and H)** (*29*). These results indicate that the Loaded-Enc sample contained both the encapsulin and EncFtn cargo proteins, whilst Empty-Enc has only the encapsulin protein.

Ferroxidase assays were performed on the samples to confirm the ability of the EncFtn loaded encapsulin nanocompartment to convert Fe(II) to Fe(III) (**Figure 1D**). This result is consistent with our previous observations for the *Rhodospirillum rubrum* Enc:EncFtn encapsulin complex(*11*). The empty encapsulin, which lacks the EncFtn cargo is enzymatically inactive.

Taken together, these data demonstrate that functionally active EncFtn has been successfully loaded into the encapsulin nanocompartment during expression in the heterologous *E. coli* host.

### The cryo-EM structure of the Loaded-Enc nanocompartment

Motivated by the apparent interior density in the Loaded-Enc sample, we performed single particle cryo-EM on the Loaded-Enc nanocompartment complex (**Table S2**). Consistent with previously published X-ray crystallographic and cryo-EM derived encapsulin models, an initial reconstruction was produced using Relion3.1 (*30*) with imposed I1 symmetry (**Figure 2, Figure S3, Figure S4** and **Table S3**) (*8, 9, 12, 27, 31*). This resulted in a reconstruction with a global resolution of 2.5 Å as determined by the gold-standard Fourier shell correlation at 0.143 (GS-FSC) (**Figure S3D**). The reconstruction displays a *T* = 1 icosahedral arrangement of sixty encapsulin monomers, with clearly resolved secondary structure elements. Small pores are visible in the shell at, or close to, the 2-, 3-, and 5-fold symmetry axes where protein monomers interact with each other (**Figure 2A**). Regions of the reconstruction around the icosahedral five-fold axes displayed a lower resolution than the other regions of the structure in a local resolution map (**Figure 2B**). This is consistent with observations of local resolution maps for the *Quasibacillus thermotolerans* encapsulin reconstruction(*12*). The monomer of the encapsulin nanocompartment from the reconstruction displays a HK97-fold typical for encapsulins (**Figure 2C**), with the position of the E-loop broadly determining the topology of the encapsulin nanocage. In the *T* = 1 Family I encapsulins this loop is at 45° to the boundary between the P- and A-domains (**Figure 2C**); whereas in the *T = 3* HK97, and *T = 3* and *T = 4* <u>encapsulins, it is found in close apposition to the A-domain. The recently described family II encapsulin from</u> *Synechococcus elongatus*(*31*) has an E-loop topology consistent with the *T = 3* and *T = 4* encapsulins, but adopts a *<u>T = 1</u>* <u>icosahedral nanocage.</u> The shift in E-loop position relative to other *T = 1* encapsulins is a result of an extended N-terminal region, that extends beneath the loop, this forces the pentameric vertices into a raised morphology consistent with the larger encapsulin nanocages.

**Figure 2:**
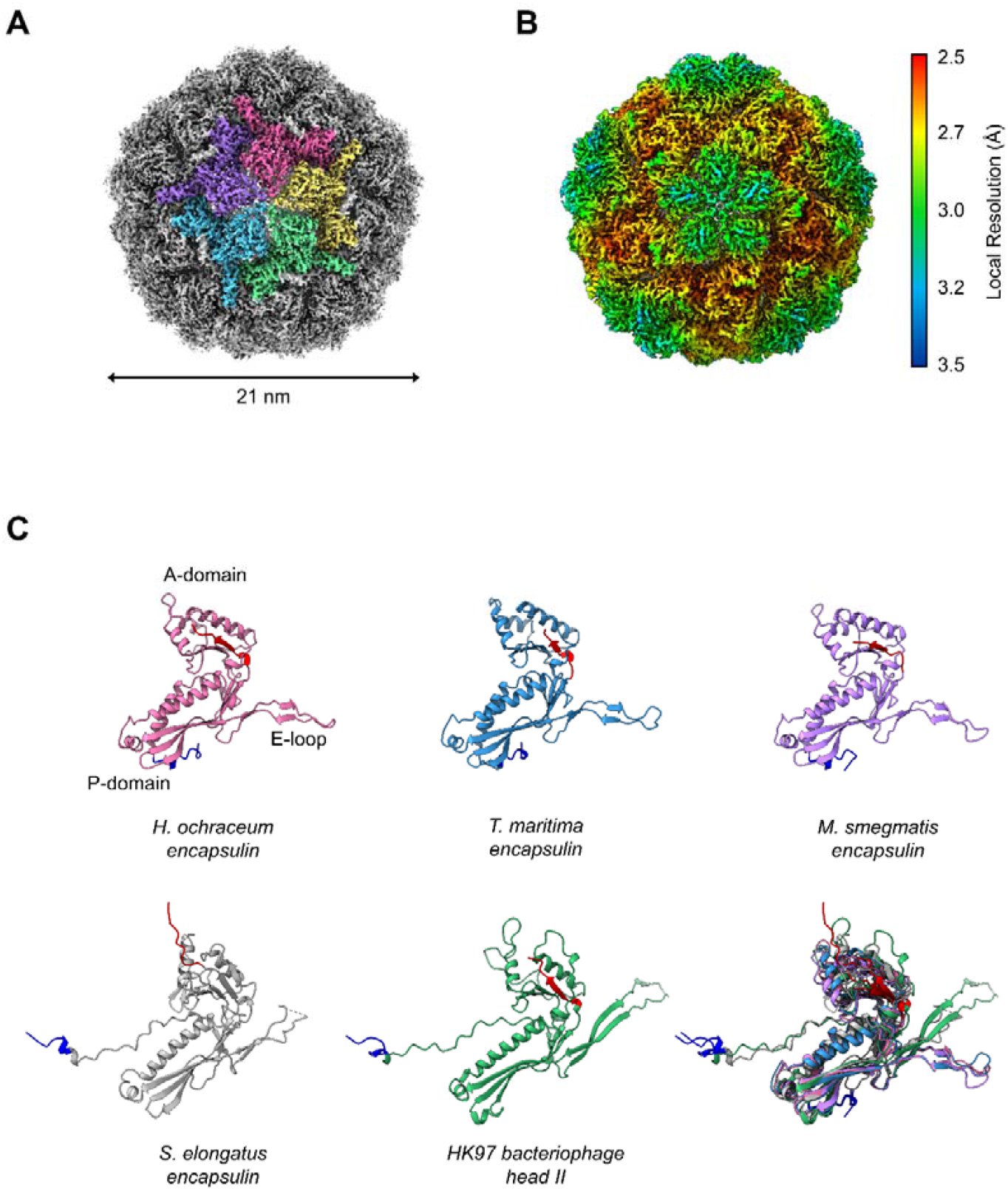
Architecture of the *H. ochraceum* encapsulin nanocompartment shell. Visualization of the electronic potential map of the *H. ochraceum* encapsulin from an icosahedrally averaged single particle reconstruction. **A**: The exterior of the encapsulin shell visualized at 2.4 Å resolution. Five subunits of the encapsulin nanocompartment shell have been colored to highlight the 5-fold axis. **B:** Icosahedral EM map of Loaded-Enc sharpened by local resolution estimate and colored by local resolution. The estimated resolution varies across the exterior of the encapsulin nanocompartment with the lowest resolution at the 5-fold pores. Color key of resolution mapping is shown on the right-hand side of the figure. **C**: The shared phage-like fold in the HK97 bacteriophage capsid and encapsulin proteins. Monomeric subunit of the *H. ochraceum* encapsulin protein modelled from our reconstruction are shown (pink), with comparisons to other *T=1* encapsulins from: *T. maritima* (blue, PDB ID:3DKT); *M. smegmatis* (lilac, PDB ID: 7BOJ); and *S. elongatu*s (grey, PDB ID: 6×8M). The HK97 bacteriophage Head II *T*=7 monomer (green, PDB ID: 2FT1) is also shown.. The N-terminus of each monomer is highlighted in blue and the C-termius in red. **Lower right:** Overlay comparison of the encapsulin monomers showing similar A- and P-domains orientations.

### The encapsulin nanocompartment recruits four EncFtn decamers to its lumen

With the imposition of I1 symmetry on the reconstruction, the EncFtn cargo is not visible within the lumen. This suggests that the organization of the EncFtn protein within the encapsulin shell does not conform to icosahedral symmetry and is rotationally averaged through our symmetry-imposed processing. To gain insight into the structural relationship between the encapsulin shell and its EncFtn cargo protein, a reconstruction was produced with no imposed symmetry averaging (**Figure 3 and Figure S5**). Separation of the dataset into five 3D classes revealed a highly populated class with amorphous density in the interior **(Figure S4, panel 5B**), and two other main classes, both containing four distinct densities, arranged in a similar tetrahedral fashion within the encapsulin shell. To obtain the clearest reconstruction of both the encapsulin nanocompartment and EncFtn cargoes we took only one of these latter classes forward for full 3D refinement in Relion3.1 to produce a final C1 reconstruction with a resolution of 3.7 Å at the 0.143 FSC threshold (**Figure 3A and Figure S5A**). Interestingly, the EM map revealed four distinct densities within the encapsulin nanocompartment lumen, which are consistent in size and shape with four EncFtn decamers (**Figure 3B** and **Figure S5B**).

**Figure 3:**
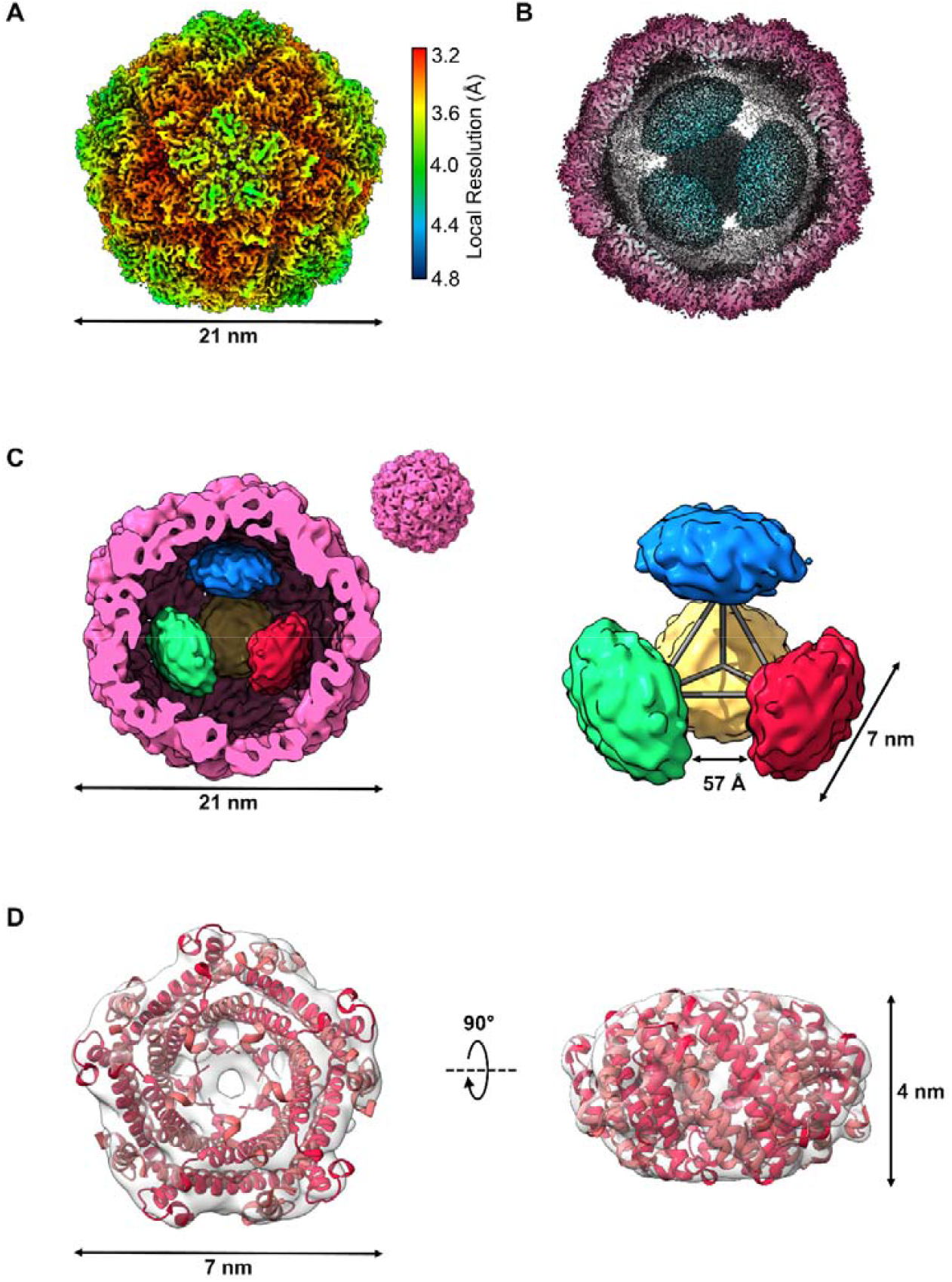
Asymmetric reconstruction of the *H. ochraceum* encapsulin complex reveals a tetrahedral arrangement of EncFtn within the encapsulin nanocompartment. **A**: Electronic potential map of the asymmetric reconstruction of the *H. ochraceum* Enc:EncFtn complex. The map is colored by local resolution with the color key shown on the right side of **A. B:** Radially colored cryo-EM derived map of the Loaded-Enc nanocompartment displaying the interior EncFtn (cyan). **C**: Gaussian smoothed C1 map showing the four discrete EncFtn densities (red, green, yellow, and blue), consistent with the size of decameric EncFtn complexes, within the encapsulin nanocompartment (pink). The four EncFtn are in a tetrahedral arrangement highlighted by grey lines connecting their centers of mass. The average distance between each EncFtn decamer is 57 Å. **D**: EM Map of a single EncFtn density following symmetry expansion and local refinement in CryoSPARC. The *H. ochraceum* EncFtn decamer crystal structure (PDB ID: 5N5F, red) is docked into the density.

A local resolution map calculated for the asymmetric reconstruction indicates a degree of flexibility at the pentameric pores compared to the trimeric pores of the encapsulin shell (**Figure 3A**). The interior of the nanocompartment shows a significant falloff in resolution from the inner face of the encapsulin shell to the EncFtn densities. This is consistent with the tethering of the EncFtn to the encapsulin nanocompartment via its localization sequence, with some degree of conformational freedom of the EncFtn decamer with respect to the encapsulin shell.

The EncFtn decamers are located approximately 3 nm away from the encapsulin interior wall, which corresponds to the linker region between the main EncFtn domain and the localization sequence on the EncFtn C-terminus (**Figure S5C**). The extended localization sequence of the EncFtn protein acts to offset it from the inner face of the encapsulin shell, an observation consistent with previous reports of the IMEF encapsulin complex from *Q. thermotolerans*(*12*). Due to the dynamic nature of the EncFtn within the encapsulin it was not possible to trace the path of the localization sequence to its binding site. Interestingly, the four EncFtn decamers are in a tetrahedral arrangement within the encapsulin nanocompartment, with the five-fold axes of the EncFtn decamers aligned to the three-fold tetrahedral axes (**Figure 3C and Figure S5C**). This results in a double symmetry mismatch between the icosahedral shell and the EncFtn decamers in the complex. A concurrent structural study of the *Thermotoga maritima* encapsulin complex(*32*) revealed five EncFtn decamers within the encapsulin shell, with each decamer found in approximation to a pentameric vertex. The overall arrangement of the five EncFtn complexes within the nanocompartment is incompatible with the formation of a regular platonic solid and results in a symmetry breaking arrangement of the EncFtn within the encapsuling nanocompartment. To enhance the resolution of the individual EncFtn density, symmetry expansion and local refinement focused on a single EncFtn density was performed in CryoSPARC. This resulted in an EM map into which the *H. ochraceum* EncFtn crystal structure (PDB ID: 5N5F)(*16*) could be docked (**Figure 3D**).

Analysis of the relationship between the EncFtn decamers and the inner face of the *H. ochraceum* encapsulin shell reveals distinct EncFtn environments (**Figure S6**). The first EncFtn environment is shared by two EncFtn decamers and is in line with the five-fold pore of the encapsulin nanocompartment (EncFtn 1 and 2 in **Figure S6**). The shared symmetry of the Enc nanocompartment five-fold pores and of the EncFtn D5 annular structure in these positions is consistent with our previously proposed hypothesis for the Enc:EncFtn relationship(*11*) and is also found in the *T. maritima* encapsulin. However, the symmetry-breaking tetrahedral arrangement of the EncFtn decamers in the *H. ochraceum* encapsulin creates a second distinct environment shared by the remaining two EncFtn decamers, where they are offset between five-fold and three-fold axes of the icosahedral encapsulin shell (EncFtn 3 and 4 in **Figure S6**). With the proposed route of iron entry through the 5-fold pores of the encapsulin shell, the EncFtn decamers in proximity to the pores may have more favorable substrate access than those found in the alternative positions. Given the fast mass-transport of substrates across the shell(*33*), this symmetry breaking arrangement of the EncFtn decamers is unlikely to have significant functional consequences in terms of the speed of iron oxidation in the different EncFtn environments.

### Structural dynamics in the pentameric vertices of the encapsulin shell

To further investigate the apparent conformational flexibility of the encapsulin shell at the pentameric vertices in the reconstructions, we analyzed the full data set in CryoSPARC to take advantage of recent advances in 3D variability analysis (3DVA) of single-particle datasets(*34*). A C1 refinement calculated with 333,945 particles performed in CryoSPARC (**Table S4**) was subjected to 3DVA with three variability components. The three components resolve to show the five-fold pores irising between ‘open’ and ‘closed’ states. The three components primarily differ in which pores are open/closed and show no other major differences. Adjacent five-fold pores show a mixture of correlated and anti-correlated opening (**Figure 4, Figure S7** and **Supplementary Movies 1-3**). This movement gives the impression of a pumping motion, which may enhance the transfer of substrates between the exterior and interior lumen of the nanocompartment.

**Figure 4:**
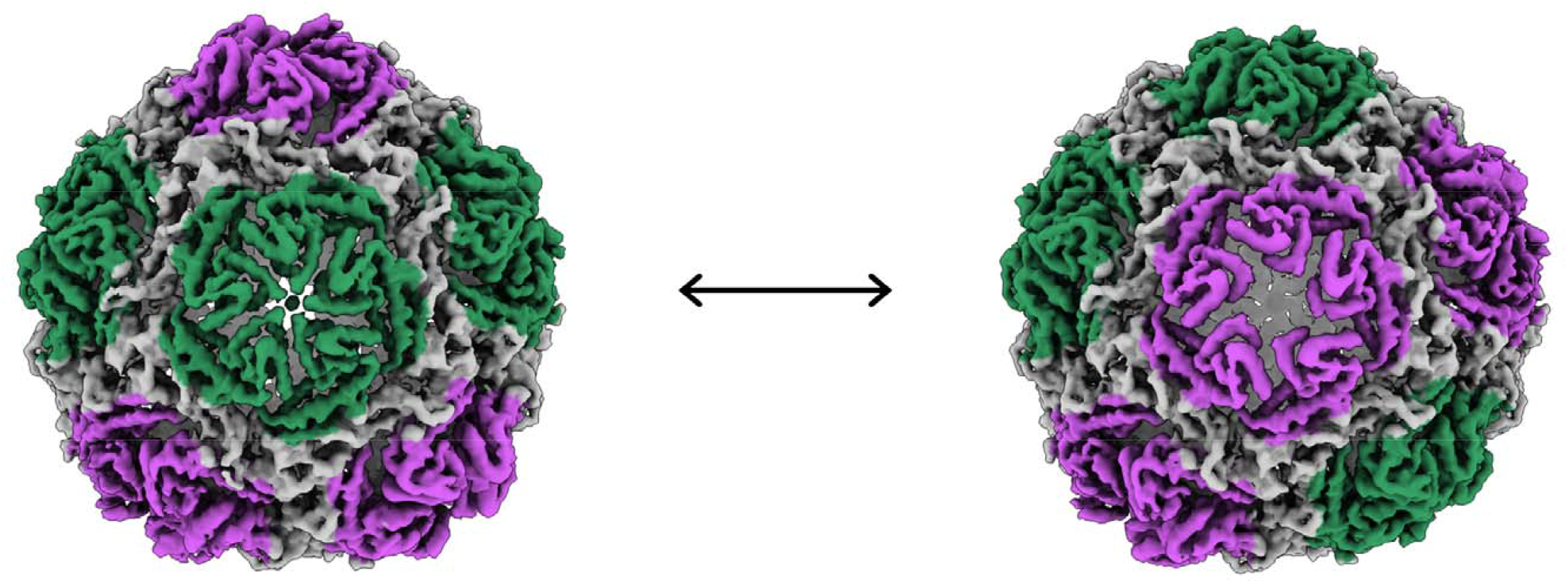
3D variability analysis of Loaded-Enc. The first and last frames from the trajectory of component 0 from the 3DVA are shown as density maps centered on a single five-fold pore. Pores across the encapsulin nanocompartment are seen to adopt conformations consistent with both ‘closed’ (green) ‘open’ states (purple) and move between these along the trajectory. Different pores adopt this dynamic in the two other components calculated.

Within the encapsulin nanocompartment, the EncFtn densities show some uncoordinated lateral motions, but their position relative to the inner face of the encapsulin does not change. This is because they are tethered by their localization sequences to the P-domain of the encapsulin, which does not show any significant displacement in any of the variability components.

To capture the extreme ‘open’ and ‘closed conformations of the five-fold pores we performed symmetry expansion on the I1 particle set in Relion3.1, followed by masked 3D-classification without alignment centered on the vertex. Of the five conformations observed, the most extreme of these were subjected to 3D refinement with local searches (**Figure 5, Figure S8**, and **Figure S9**). This resulted in an ‘open’ pentamer conformation of 2.4 Å resolution and a ‘closed’ conformation of 2.3 Å, allowing for fitting of residue side chains (**Figure S7**).

**Figure 5:**
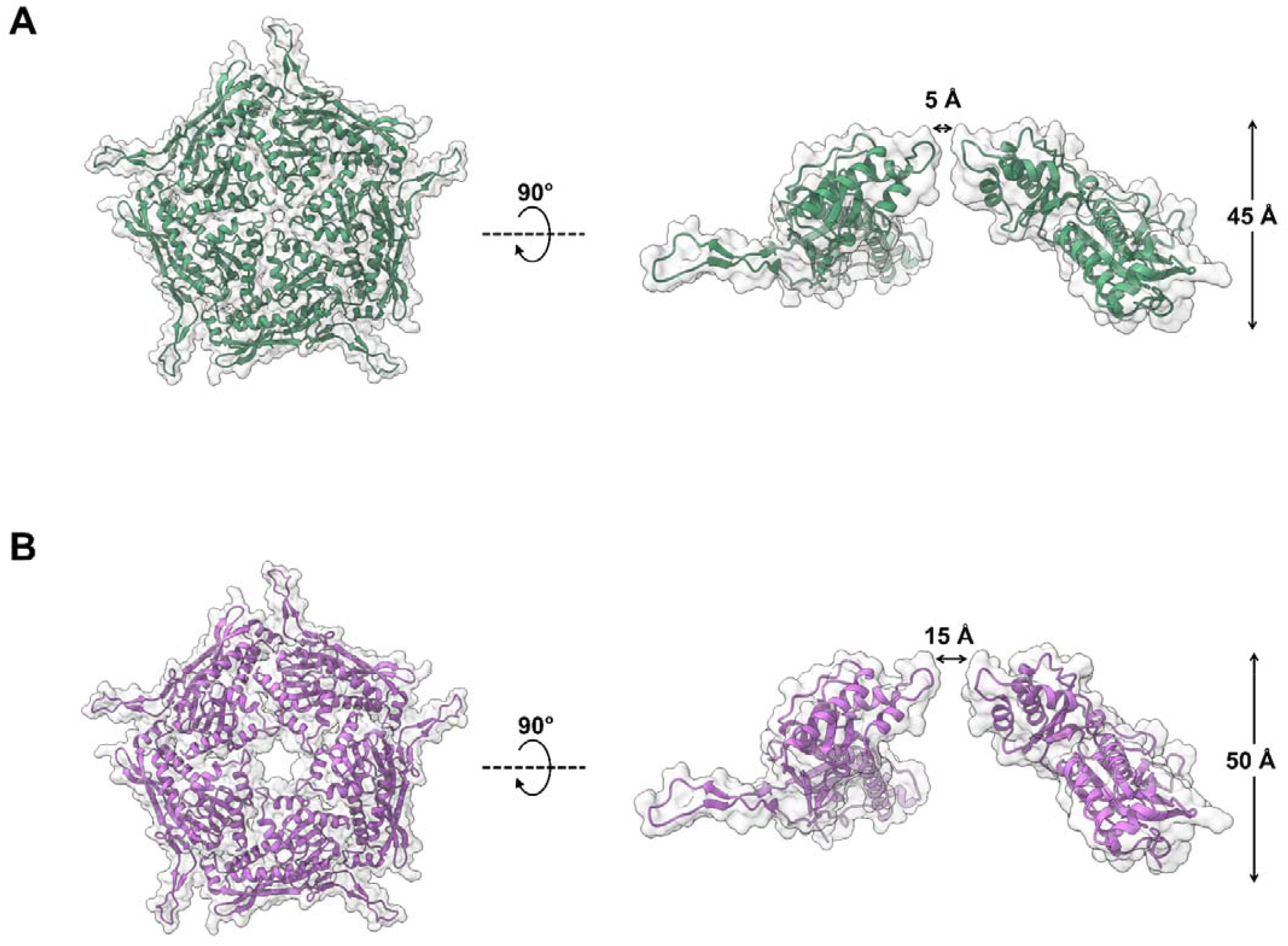
Open and closed conformations of the dynamic 5-fold pore of the *H. ochraceum* encapsulin shell. Masked 3D refinements centered around the 5-fold pore of the icosahedral reconstruction were performed after symmetry expansion of the asymmetric units. The ‘closed’ (**A**, green) and ‘open’ (**B**, purple) conformations of the Loaded-Enc shell pentamer are represented with transparent solvent accessible surface rendered over cartoon secondary structure. Both ‘closed’ and ‘open’ pentamers are shown in top-down (left-hand side) and side-on views (right-hand side). The closed conformation has a 5-fold pore diameter of 5 Å, whilst the equivalent diameter of the open conformation is 15 Å. To allow for the widening of the pore, the open pentamer protrudes from the nanocompartment by 5 Å.

The ‘open’ conformation has a five-fold pore with an aperture diameter of approximately 15 Å, while in the ‘closed’ conformation the aperture is reduced to 5 Å diameter. To understand the structural changes taking place in the transition between these conformations, an atomic model of the encapsulin protein was refined against both maps (**Table S5**). The two models show a significant movement in the A-domain, with the A-domain pivoting at the loops connecting it to the P-domain (residues 121 – 131 and 201-222) (**Figure S10A/B**), opening the pore like an iris (**Figure 5**). In the open conformation, the pore loop region (residues 182 - 189) is not well defined in the density; while it is tightly locked in the closed conformation, with Asp186 forming the outer boundary of the pore and Tyr188 and Lys192 forming the inner bounds (**Figure S10C**). The tyrosine is well conserved among the family 1 *T=*1 encapsulins, while the lysine is substituted for a glutamine in the *R. rubrum* encapsulin (**Figure S11**). The family 2 *T*=1 encapsulin from *Synechococcus elongatus* has a five-residue sequence insertion in this region, which forms an extended linker between secondary structure elements, rather than a distinct loop within the pore.

Additionally, our focused refinements of the pentameric subunits allowed us to build and sequence the _117_GSLGIGSLR_125_ peptide from the EncFtn protein (**Figure 6**). This region of the localization sequence forms a network of hydrophobic interactions with the inner face of the P-domain of a single encapsulin monomer, with further stabilization by several water-mediated backbone contacts. The core GxLGIxxL motif found in this region of the localization sequence is conserved between the *H. ochraceum* EncFtn and other proteins in the family and is observed in the crystal structure of the *T. maritima* encapsulin(*9*).

**Figure 6:**
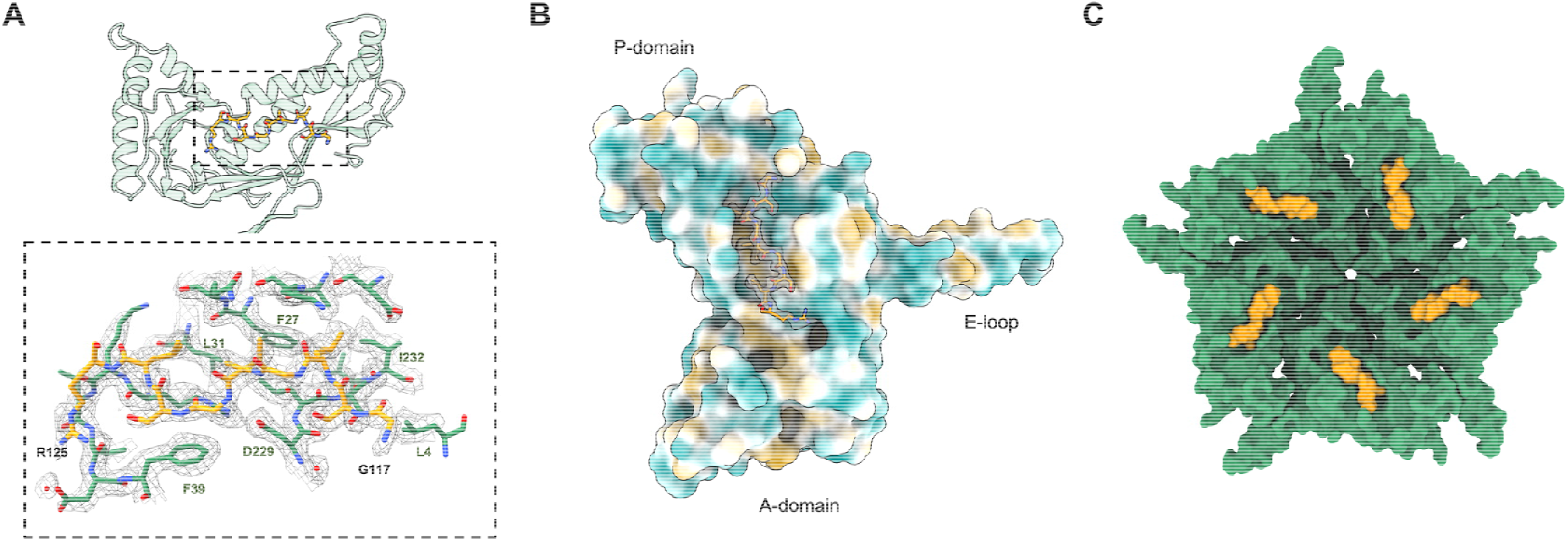
The closed conformation of the *H. ochraceum* five-fold pore allows docking and sequencing of the EncFtn localization sequence to the interior face of the encapsulin monomer. **A**: Binding of localization sequence residues from the EncFtn to the interior wall of the encapsulin nanocompartment monomer (green residues and transparent cartoon) and the localization sequence (yellow sticks). Lower panel: Hydrophobic residues from the interior face of the encapsulin form the binding pocket for the localization sequence. The first and last residues modelled for the LS have been labelled in black, and key residues from encapsulin have been labelled in green. Modelled water molecules are shown as red spheres. **B:** The spatial relationship between an encapsulin monomer and the localization sequence (shown in gold with its EM map density as in **A**). The Enc monomer has been colored by molecular lipophilicity potential (*54*) which ranges from dark cyan (corresponding to the most hydrophilic) to white to dark gold (most lipophilic). This highlights the hydrophobic pocket on the interior of the encapsulin nanocompartment where the localization sequence binds. **C**: The ‘closed’ conformation pentamer (green) with a localization sequence (yellow) shown on each monomer.

In the *H. ochraceum* encapsulin the five-fold pore has a negative charge on the exterior of the encapsulin shell and positive charge on the interior in both the open and closed conformations (**Figure S12**). No changes are observed in the structure, or charge of the 3- and 2-fold pores (**Figure S12**). The closed conformation of the 5-fold pore is consistent with observations from the crystal structure of the *T. maritima* encapsulin(*9*) and high resolution cryo-EM structures of other encapsulins(*12, 31, 32*). However, this is the first time that an ‘open’ pore-conformation has been observed in an encapsulin protein.

This observation has important implications for efforts to engineer the pores of encapsulin nanocages. Efforts to widen the five-fold pores have demonstrated an increase in mass-transport of model substrates and ions across the encapsulin shell(*33, 35*); however, the observed changes in pore diameters are limited to a maximum of around 10 Å in the *T. maritima* encapsulin(*35*). Interestingly, the encapsulin shell appears to interact strongly with Tb^3+^ ions in molecular dynamics simulations, this is consistent with our previous observations of iron binding by the *R. rubrum* encapsulin(*11*) and highlights a possible cooperative role for substrate channelling to the EncFtn cargoes.

Our results suggest that the flexibility of the A-domain should also be considered when engineering the pore structure for enhanced transport across the encapsulin shell. The loops connecting the A-domain to the P-domain present ideal sites for targeted mutagenesis to stiffen, or lock, the position of the A-domain and thus constrain the five-fold pore loop to engineer enhanced substrate selectivity into these systems.

### Dynamics of the 5-fold encapsulin pore through hydrogen/deuterium exchange mass spectrometry

To further investigate the dynamic nature of the five-fold pore of the encapsulin shell and the docking of the EncFtn localization sequence to the interior of the encapsulin nanocompartment, we performed hydrogen/deuterium exchange mass spectrometry (HDX-MS) on both Empty-Enc and Loaded-Enc nanocompartments. The extent of backbone-amide hydrogen exchange was determined at seven time points (0 seconds, 10 seconds, 30 seconds, 5 minutes, 30 minutes, 4 hours, and 24 hours). By calculating the rate of hydrogen exchange throughout the protein, regions that differ in solvent exposure and/or dynamics can be detected.

HDX-MS analysis of the encapsulin nanocompartment resulted in 40 pepsin peptides, which constituted a protein sequence coverage of 85%, with peptide redundancy of 2.28 (**Table S6, Table S7** and **Figure S13**). The encapsulin nanocompartment displayed variable exchange rates throughout the protein sequence and regions of the protein displaying elevated H/D exchange rates were evident. Overlaying these local H/D exchange rates onto the cryo-EM reconstruction revealed that the regions of highest exchange were located around pentameric vertices. (**Figure 7**). This was most notable with the peptide spanning the region between amino acids 180-196, which includes the five-fold pore loop (**Figure S14**). In contrast, lower rates of HDX are observed at the 2-fold interface and the potential 3-fold pore (**Figure 7**). These findings agree with our cryo-EM structural analyses and support the proposed conformational flexibility at the 5-fold pore.

**Figure 7:**
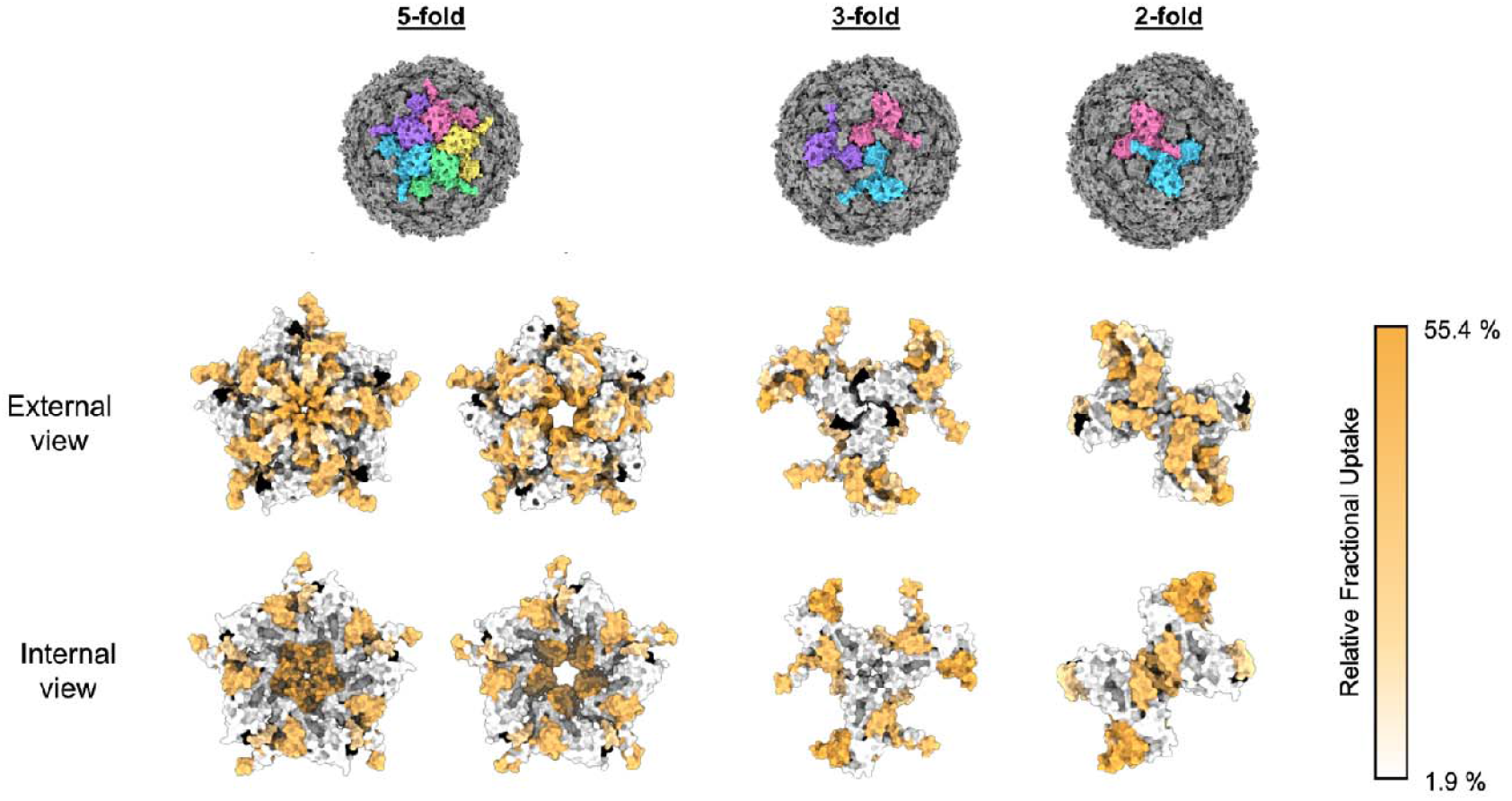
HDX-MS fractional uptake mapped to the symmetry axes of the icosahedral reconstruction of the *H. ochraceum* encapsulin complex. From top to bottom: The top row depicts molecular surface models of the *H. ochraceum* encapsulin in three orientations to show the 5-fold, 3-fold and 2-fold symmetry axes. Monomers have been colored individually (pink, yellow, green, blue, and purple) to highlight the symmetry axes. The final two rows display the relative fractional uptake of each peptide from HDX experiments displayed on the 5-fold, 3-fold and 2-fold pores of the encapsulin nanocompartment. Peptide fractional uptake is show on a white-to-orange color scale with a color key shown on the right-hand side of the figure. Areas colored black correspond to no peptide coverage.

Comparison of the H/D exchange rates of Empty-Enc and Loaded-Enc revealed similar exchange profiles throughout the encapsulin protein sequence, suggesting that cargo loading has little effect on the overall architecture and dynamics of the assembled nanocompartment shell (**Table S6, Table S7**, and **Figure S15**). However, after prolonged exchange times (4 hours), the Loaded-Enc exhibited areas with a modest reduction in exchange when compared to the Empty-Enc. Notably, several peptides in the N-terminal region displayed reduced exchange rates in Loaded-Enc; for example, the peptide covering amino acids 21-37 displayed almost twelve percent reduction. Mapping the position of this region onto our encapsulin reconstruction highlights that this peptide is located on the interior face of the nanocompartment and included the proposed binding site for the localization sequence of EncFtn (**Figure 7**). A reduction in exchange across in this region is likely a consequence of shielding by the engaged EncFtn localization sequences.

## Discussion

Our cryo-EM reconstruction of an Enc:EncFtn nanocompartment complex reveals key areas of divergence from a true icosahedral complex with important functional consequences. The asymmetric reconstruction showed that the encapsulin nanocompartment sequesters four decamers of EncFtn within its lumen. In our recombinant system, with EncFtn produced in excess, this likely represents a maximum loading capacity for the Enc:EncFtn nanocompartments. The symmetry breaking tetrameric arrangement of the EncFtn decamers within the encapsulin shell leads to two distinct environments for EncFtn, with two decamers aligned at the five-fold symmetry axes, and the remaining two residing between three- and five-fold axes (**Figure S6**). The concurrent observation of five EncFtn decamers within the *T. maritima* encapsulin nanocompartment, aligned close to the icosahedral five-fold symmetry axes of the encapsulin shell, highlights differences in cargo loading between encapsulins from mesophilic and thermophilic bacteria(*32*). The enhanced cargo loading seen in the *T. maritima* encapsulin may be a consequence of a more compact and rigid structure adopted by thermophilic proteins, as seen in comparisons of the crystal structures of the *H. ochraceum* and *Pyrococcus furiosus* EncFtn proteins(*16*).

The interior volume of the *T=1* encapsulin lumen is around 4000 nm^3^, while an EncFtn decamer is only 120 nm^2^; therefore, the additional cargo seen in the *T. maritima* encapsulin should not significantly impact the iron-storage potential. The order of magnitude discrepancy in the iron-loading capacity measured for the *R. rubrum*(*11*) and *T. maritima*(*32*) encapsulins is likely a result of differences in experimental conditions in different laboratories.

While the volume occupied by the EncFtn decamers represents less than 15 % of the total lumen of the encapsulin nanocompartments, the consistent observation of a gap between the encapsulin shell and encapsulin cargo proteins implies that the loading and capacity of encapsulins is limited by broader steric effects. Further steric effects would also be found from the unengaged localization sequences at the core of the nanocompartment. These effects have implications for efforts to target heterologous proteins to the encapsulin nanocage; effectively setting a limit on the volume of protein that can be accommodated within, which is much lower than the total volume of the lumen of the nanocage.

Our observations have functional implications for the oxidation and storage of iron within the Enc:EncFtn nanocompartment. The EncFtn decamers are in non-equivalent positions, and thus have different relationships to the pores of the encapsulin shell. Therefore, if the pores limit the diffusion of substrates, the EncFtn decamers would be subjected to different chemical environments. Furthermore, both the engaged and unengaged localization sequences present a ‘soft’ steric barrier to the diffusion of substrates. It is notable that the ferroxidase activity of the Enc:EncFtn complex is significantly higher than the isolated EncFtn protein(*36*). The dynamic behavior of the encapsulin five-fold pores, which is apparent in our 3DVA movies, may increase EncFtn activity by enhancing diffusion of substrates into the lumen of the nanocompartment. While it is not possible to make mechanistic conclusions from our model, the complex interactions with the components of the encapsulin nanocompartment clearly enhance the iron oxidation activity of the EncFtn protein.

Our data were collected on iron-free apo-Enc:EncFtn complexes, and thus it is not possible to infer the nature of the iron mineralization pathway within the encapsulin nanocage. Further careful work must be performed to titrate iron into the complex prior to structural analysis to gain insight into the flow of metal ions from the exterior to the interior of the encapsulin and to determine if metalation influences the conformational flexibility of the EncFtn within the encapsulin nanocage, as we have demonstrated for isolated EncFtn proteins(*11, 16, 36*). Finally, the nature of the iron mineral and its localization within the encapsulin nanocage is still to be determined.

These cryo-EM and HDX-MS data illustrating a highly dynamic five-fold pore in the encapsulin shell, have major implications for efforts to engineer recombinant encapsulins for improved access for both native and non-native substrates. The limitations of previously published studies, where the five-fold pore is modified for altered substrate access can be explained by a highly dynamic pore structure. Our work suggests new hypotheses for engineering pore selectivity, through modifications to the hinge regions between the P- and A-domains, which are responsible for the opening of the pore.

## Materials and Methods

### Experimental Design

The objective of this study was to understand the structural relationship between encapsulins and their EncFtn cargo using cryo-EM to determine the complex structure and ferroxidase assays for validation of the complex activity. HDX was utilized to establish differences between solvent accessibility of empty and loaded encapsulins and garner insight into the impact of cargo on the interior of the encapsulin nanocompartment.

### Cloning of encapsulin expression constructs

The *Haliangium ochraceum* encapsulin and encapsulated ferritin protein expression constructs were based on the Hoch_3836 and Hoch_3837 genes downloaded from www.kegg.jp and were codon optimized for expression in *Escherichia coli* and synthesized as CIDAR MoClo compatible gBlocks by Integrated DNA Technologies (IDT) (**Table S8**). The gBlocks were assembled into a Level 0 CIDAR MoClo storage vector(*37*), DVA_CD, for subsequent use. The coding sequences for the encapsulin and the EncFtn were assembled into expression cassettes in the level 1 backbones DVK_AE and DVK_EF respectively, each with T7 promoter and transcription terminator parts. The resulting expression cassettes were then combined into the DVA_AF backbone to produce a co-expression plasmid. All assembled plasmids were sequence verified by Sanger sequencing by Eurofins Genomics. The protein sequences for each construct are listed in **Table S9**.

### Protein expression

The Empty-Enc and Loaded-Enc expression plasmids were transformed into *E. coli* BL21(DE3) cells and grown overnight at 37 °C on LB-agar plates containing appropriate selection antibiotics (kanamycin for Empty-Enc and ampicillin for Loaded-Enc). A single colony of cells was added to 1 L of autoinduction media(*38*) supplemented (**Table S10)** with appropriate antibiotic and grown for 38 hours at 37 °C with shaking at 200 rpm. Cells were harvested by centrifugation at 12,000 × *g*.

### Encapsulin nanocompartment purification

*E. coli* cell pellets expressing the empty-Enc and loaded-Enc constructs were resuspended in 10 × v/w of lysis buffer (20 mM HEPES, pH 8; 2 mM MgCl_2_; 1 mg/mL lysozyme, and benzonase, 12.5 – 25 units/mL). Cells were lysed by sonication whilst on ice; sonication was carried out in six 1-minute cycles (30 seconds sonication, 30 seconds rest). The lysate was clarified by centrifugation at 20,000 × g for 1 hour, 4 °C.

The supernatant from cell lysis was heated to 85 °C for 10 minutes in a water bath and transferred to a 4 °C ice bath for 10 minutes. The supernatant was then collected after centrifugation at 10,000 × g for 1 hour.

Anion exchange chromatography of the clarified supernatant was performed using a 1 mL HiTrap Q Sepharose FF column from Cytiva on an ÄKTA™ start. The column and ÄKTA™ start system were equilibrated with QA buffer (20 mM HEPES, pH 8.0) and the protein sample was loaded. Unbound proteins were removed by washing with QA buffer. Bound proteins were eluted by QB buffer (20 mM HEPES, pH 8.0, 1 M NaCl) over a linear gradient of 0-100% QB over 15 column volumes. Flow-through fractions containing the sample were subjected to SDS-PAGE to identify those containing the protein of interest. These fractions were pooled and concentrated using centrifugal concentrators with a 30 kDa nominal molecular weight cut off (Vivaspin).

Pooled and concentrated samples from the anion exchange step were loaded on a size exclusion chromatography column (Sephacryl 400, Cytiva) equilibrated with SEC buffer (20 mM HEPES, pH 8.0, 150 mM NaCl). Fractions eluting from the column containing the desired protein, as identified by SDS-PAGE were pooled and concentrated as above. Protein aliquots were flash cooled in liquid nitrogen and stored at −80 °C (**Figure S1Ai, Aii, Bi and Bii**). (*11, 16*).

Following purification, aliquots of Empty-Enc and Loaded-Enc were subjected to a second round of polishing size exclusion chromatography using an Sephacryl 400 column (Cytiva) equilibrated with SEC buffer (20 mM HEPES, pH 8.0, 150 mM NaCl) (**Figure 1B**). (Experimental data for the size-exclusion chromatography runs are available at: doi: <u>10.6084/m9.figshare.16698106</u>)

### Negative stain TEM

Purified encapsulin nanocompartments were initially imaged by negative stain TEM. Continuous carbon/formvar coated copper grids (200 mesh) were glow-discharged for 30 seconds using a Pelco glow discharge system. 5 μL Enc was pipetted onto the glow-discharged grids and excess liquid was removed after 30 seconds with Whatman filter paper (grade 1, diameter 24.0 cm). The grids were washed with distilled water three times, followed by staining with 2 % uranyl acetate for 5 seconds. Grids were left to air dry and then imaged with a JEOL JEM-1400 transmission electron microscope. Images were collected with a Gatan CCD OneView camera and analyzed using FIJI(*39*). (Raw negative stain images are available at: doi: <u>10.6084/m9.figshare.16698088</u>)

### Ferroxidase activity assay

The enzymatic activity of Empty-Enc and Loaded-Enc were assessed by ferroxidase assay, as previously described (Piergentili, 2020). Fe(II) samples were prepared by dissolving FeSO_4_.7H_2_O in HCl 0.1 % (v/v) under anaerobic conditions. Protein samples were diluted anaerobically in Buffer GF (20 mM HEPES, pH 8.0, 150 mM NaCl) to a final encapsulin monomer concentration of 9 μM to allow comparison between experiments.

Iron and protein aliquots were added aerobically to a quartz cuvette (Hellma) resulting in a final concentration of 100 μM iron and 15 μM (Loaded-Enc), or 9 μM (Empty-Enc). The cuvette was placed in a UV-visible spectrophotometer (PerkinElmer Lambda 35) and the reaction sample was incubated at 21°C for 50 s to stabilise. Absorbance at 315 nm was then recorded every second for 1450 s using the Time-Drive software. A control experiment was conducted by monitoring the background oxidation by atmospheric oxygen of 100 μM FeSO_4_*7H_2_O in the absence of the enzyme. Loaded-Enc experiments were carried out with three biological replicates. There were six technical replicates for batch one, two relplicates using batch two and one replicate from batch three. Means and standard deviations were calculated on the time zero-subtracted progress curves. (Experimental data for the ferroxidase assays are available at: doi: <u>10.6084/m9.figshare.16698127</u>)

### Liquid Chromatography Mass Spectrometry

LC-MS experiments were performed on a Synapt G2 Q-ToF instrument (Waters Corp., Manchester, UK) and an Acquity UPLC equipped with a reverse phase C4 Aeris Widepore 50□×□2.1Lmm HPLC column (Phenomenex, CA, USA). Mobile phases of; A= water + 0.1% formic acid, and B=acetonitrile + 0.1% formic acid were used on a ten-minute gradient from 5% B to 95% B. Samples were analyzed at ∼2 μM, and data analysis was performed using MassLynx v4.1 and MaxEnt deconvolution.

### Cryo-EM data collection and analysis

#### Sample vitrification

Holey grids (gold, 200 mesh, r 2/2 by Quantifoil) were glow-discharged for 30 seconds using a Pelco glow discharge system. The grids were then mounted into a FEI vitrobot and 4 μL of encapsulin sample (3 mg/mL) was applied. Grids were then blotted (100% humidity, 8 °C, blot force −5, wait time 10 seconds and blot time of 3 seconds) with Whatman filter paper (grade 1) and flash cooled in liquid ethane, cooled with liquid nitrogen.

#### Cryo-EM data collection

Cryo-EM grid screening was performed on a FEI F20 microscope equipped with a FEG electron source (200 kV) and a TVIPS F816 CMOS detector at the University of Edinburgh. The dataset used for single particle reconstruction was obtained at eBIC on a FEI Titan Krios microscope equipped with Gatan K3 camera (data collection settings are shown in **Table S2**). Alignments, grid transfer and imaging set up was performed by the eBIC local contact Dr Yun Song. (Cryo-EM movies are available in the EMPIAR database: <u>EMPIAR-1218</u>).

#### Single particle reconstruction

Processing steps for the main reconstruction were performed with the Relion3.1 software package (*30*). Super-resolution movies were binned at 2 × 2 pixels for both I1and C1 and motion corrected using MotionCor2(*40*). Defocus values of images (summed movies) were determined by CTFFIND4 (*41*) and those with poor CTF fits, or bad ice, were manually discarded. A template for autopicking was created using 2D classes from manually picked particles. Autopicked particles were extracted (using box a box size of 512 pixels and subjected to three rounds of 2D classification to remove bad particles. An initial 3D model was created from particles selected from 2D classes. 3D classes were generated both with and without icosahedral symmetry imposed (‘I1’ and ‘C1’ symmetry). The best class from each was taken forward for 3D refinement and then CTF refinement followed by further rounds of 3D refinement and Bayesian polishing. After a final round of 3D refinement, postprocessing was performed using a soft spherical mask. Local resolution estimation was performed in Relion3.1. The data processing and refinement pipeline is shown in **Figure S3** with data processing and refinement statistics in **Table S3**.

3D variability analysis was performed using CryoSPARC(*32, 41*). Super-resolution movies were patch motion corrected and patch CTF estimated. Initally, particles were picking using blob picker (250-300 Å) and these particles were subjected to 2D classification followed by a round of ab initio model generation to create an autopick template. Successive rounds of rounds of 2D classification were performed on the autopicked particles until less than 5% of the particles were rejected. Five volumes were created by the ab initio model generation algorithm, and these were heterogeneously refined. The volume with the most particles (99.4 %) was homogeneously refined and subsequently subjected to 3D variability analysis(*34*). The homogenously refined particles were also used to enhance the resolution of the EncFtn cargo. Particles were subjected to symmetry expansion, and then an individual EncFtn was masked and subjected to local refinement. The data processing and refinement performed in CryoSPARC pipeline is shown in **Figure S4** with data processing and refinement statistics in **Table S4**. Movies were created with ChimeraX(*43*).

Motivated by the dynamic behaviour of the 5-fold pores of the encapsulin shell, the 5-fold pore pentamer was subjected to symmetry expansion and focused classification. The I1 particle set was expanded using the sym_expand job in Relion3.1 and masked 3D classification without alignment was performed focused on the five-fold symmetry axis. Of the five classes produced the two most highly populated and distinct were taken forward for masked 3D refinement with local searches only, these two classes represent the ‘open’ and ‘closed’ conformation of the five-fold pore. The mask used in these steps was produced in Chimera using the molmap command on a docked model of a pentamer of the encapsulin protein. To ensure the box-size and pixel-size of the mask were correct, they were resampled onto the icosahedral map using the vop resample command.

#### Model building and refinement

An initial homology model of the encapsulin nanocompartment monomer was generated using Phyre 2.0(*44*) based on the *T. maritima* structure. This was docked into the open and closed maps using ChimeraX(*45*) and expanded to a full pentamer model. The model was then fit to the map through an iterative process of automated model refinement with phenix real-space refinement(*46*) and manual model building in Coot(*47*), waters were added using phenix.douse and validated in in Coot. The resulting models and maps were validated using Molprobity(*48*), phenix.mtriage(*49*), and EM ringer(*50*) (**Table S2**). Models and maps were visualized using ChimeraX.

### Encapsulin sequence analysis

Encapsulin sequences were obtained from the Kyoto Encyclopedia of Genes and Genomes (www.kegg.jp). Sequence alignments were performed using Clustal Omega(*51*) and visualized using ESPript(*52*).

### Hydrogen/Deuterium Exchange Mass Spectrometry

Hydrogen/deuterium exchange mass spectrometry (HDX-MS) was performed on a Synapt G2 MS system coupled to an ACQUITY UPLC M-Class UPLC with the HDX manager module (Waters Corporation, Manchester, UK)(*53*). For improved reliability and precision, a custom-built Leap automated platform was utilized in all sample preparation and injections. Prior to HDX-MS analysis, three buffer solutions were prepared – (i) equilibration buffer (4.7 mM K_2_HPO_4_, 0.3 mM KH_2_PO_4_ in H_2_O), adjusted to pH 8.0 with formic acid; (ii) labelling buffer (4.7 mM K_2_HPO_4_, 0.3 mM KH_2_PO_4_ in D_2_O), adjusted to pH 8.0 with DCl and quench buffer (50 mM K_2_HPO_4_, 50 mM KH_2_PO_4_ in H_2_O) adjusted to pH 2.3 with formic acid. Protein samples were diluted in equilibration buffer to final stock concentration of 42 mM. The time course experiments consisted of 7 timepoints: T0 (0 minute; undeuterated control), T1 (20 seconds), T2 (30 seconds), T3 (2 minutes), T4 (5 minutes) and T5 (30 minutes) T6 (4 hours) and T7 (24 hours) with each timepoint being performed in triplicate. Sample preparation consisted of 5 μL protein solution, 57 μL equilibrium buffer (T0) or labelling buffer (T1-7). The final concentration of deuterium during the labelling step was 91.2 %. Exchange was allowed to proceed at 4 °C. To arrest the exchange reaction, 50 μL of quench buffer was added to this initial solution just prior to sample injection.

After injection, samples underwent proteolytic digestion on a 2.1 × 30 mm Waters Enzymate BEH pepsin column for 3 minutes at 200□μL/min. After digestion, the peptide digest was loaded on to a 2.1 × 5.0 mm Acquity BEH C18 VanGuard 1.7μm C18 Trapping column to pre-concentrate the sample for 3□minutes at 200□μL/min. Following trapping, the digests were separated through a 2.1 × 5.0 mm Acquity BEH 1.7 μm analytical column prior to MS/MS (MS^e^) analysis via the Water Synapt G2 MS system running MassLynx v4.1 software (Waters Corporation, Manchester, UK). The separation gradient was 5-95 % acetonitrile with 0.1 % formic acid over 12 minutes at 40 μL/min. Both the trapping and LC separation were performed at 1 °C to minimize back exchange. Post-processing was performed using Proteinlynx Global Server 3.0.3 and Dynamx 3.0 software to determine the average deuterium uptake for each peptide at each time point. For comparative analyses, the relative fractional uptake was determined by dividing the observed deuterium uptake by the number of available amide exchangers on each peptide.

## Acknowledgements

We acknowledge Diamond Light Source for access and support of the cryo-EM facilities at the UK’s national Electron Bio-imaging Centre (eBIC) [under proposal EM16637], funded by the Wellcome Trust, MRC and BBRSC. We thank Dr Yun Song for their assistance at eBIC with sample loading and data collection setup. This research made use of the Rocket High Performance Computing service at Newcastle University; we thank Dr Karen Bower for assistance with the use of the Rocket service. Negative stain TEM was performed in Edinburgh Electron Microscopy Lab and the Newcastle University Electron Microscopy Research Services, we thank Steve Mitchell, Tracey Davey, and Ross Laws for their technical support. Cryo-EM grid screening was performed in the Edinburgh cryo-EM facility in School of Biological Sciences at the University of Edinburgh. The cryo-EM facility was set up with funding from the Wellcome Trust (087658/Z/08/Z) and SULSA and is supported by the Wellcome Centre for Cell Biology. We thank Professor Martin Noble for generously donating GPU time to our CryoSPARC analyses.

## Data Availability

Please see Experimental Procedures and tables for links to these. All data needed to evaluate the conclusions in the paper are present in the Supplementary Materials or have been deposited in appropriate publicly available data repositories. Please see the Materials and Methods sections and experimental tables for links to these datasets.

## Author Contributions

Conceptualization: JR, JMW, DJC

Methodology: JR, ZM, TL, CP, AMBL, JMW, DJC

Validation: JR, ZM, TL, JMW, DJC

Formal analysis: JR, ZM, TL, AMBL, PJ, JMW, DJC

Investigation: JR, ZM, KG, JEB, CP, EZA

Resources: KG, FC, CLM, JEB, AMBL, EZA, MDW

Data curation: JR, TL, DJC, AMBL, JMW

Writing – original draft preparation: JR, DJC, JMW

Writing – review and editing: JR, ZM, CP, AMBL, MDW, KJW, JMW, DJC

Visualization: JR, ZM, TL, DJC, JMW

Supervision: JMW, DJC

Funding acquisition: MDW, LEH, KJW, DJC, JMW

## References

1. Y. Diekmann, J. B. Pereira-Leal, Evolution of intracellular compartmentalization. Biochem. J. 449, 319–331 (2013).

2. C. A. Kerfeld, S. Heinhorst, G. C. Cannon, Bacterial Microcompartments (2010) (available at http://www.annualreviews.org/eprint/uv9wHHypNKsEb5yrt6z8/full/10.1146/annurev.micro.112408.134211).

3. W. Bonacci, P. K. Teng, B. Afonso, H. Niederholtmeyer, P. Grob, P. A. Silver, D. F. Savage, Modularity of a carbon-fixing protein organelle. Proc. Natl. Acad. Sci. 109, 478–483 (2012).

4. C. V Iancu, D. M. Morris, Z. Dou, S. Heinhorst, C. Gordon, G. J. Jensen, NIH Public Access. 396, 105–117 (2010).

5. T. O. Yeates, C. A. Kerfeld, S. Heinhorst, G. C. Cannon, J. M. Shively, Protein-based organelles in bacteria: carboxysomes and related microcompartments. Nat. Rev. Microbiol. 6, 681–691 (2008).

6. S. C. Andrews, The Ferritin-like superfamily: Evolution of the biological iron storeman from a rubrerythrin-like ancestor. Biochim. Biophys. Acta. 1800, 691–705 (2010).

7. E. Chiancone, P. Ceci, A. Ilari, F. Ribacchi, S. Stefanini, Iron and proteins for iron storage and detoxification. BioMetals. 17, 197–202 (2004).

8. F. Akita, K. T. Chong, H. Tanaka, E. Yamashita, N. Miyazaki, Y. Nakaishi, M. Suzuki, K. Namba, Y. Ono, T. Tsukihara, A. Nakagawa, The crystal structure of a virus-like particle from the hyperthermophilic archaeon Pyrococcus furiosus provides insight into the evolution of viruses. J. Mol. Biol. 368, 1469– 83 (2007).

9. M. Sutter, D. Boehringer, S. Gutmann, S. Günther, D. Prangishvili, M. J. Loessner, K. O. Stetter, E. Weber-Ban, N. Ban, Structural basis of enzyme encapsulation into a bacterial nanocompartment. Nat. Struct. Mol. Biol. 15, 939–947 (2008).

10. T. W. Giessen, P. A. Silver, Widespread distribution of encapsulin nanocompartments reveals functional diversity. Nat. Microbiol. 2, 17029 (2017).

11. D. He, S. Hughes, S. Vanden-Hehir, A. Georgiev, K. Altenbach, E. Tarrant, C. L. Mackay, K. J. Waldron, D. J. Clarke, J. Marles-Wright, Structural characterization of encapsulated ferritin provides insight into iron storage in bacterial nanocompartments. Elife. 5, e18972 (2016).

12. T. W. Giessen, B. J. Orlando, A. A. Verdegaal, M. G. Chambers, J. Gardener, D. C. Bell, G. Birrane, M. Liao, P. A. Silver, Large protein organelles form a new iron sequestration system with high storage capacity. Elife. 8, 1–23 (2019).

13. C. A. McHugh, J. Fontana, D. Nemecek, N. Cheng, A. A. Aksyuk, J. B. Heymann, D. C. Winkler, A. S. Lam, J. S. Wall, A. C. Steven, E. Hoiczyk, A virus capsid-like nanocompartment that stores iron and protects bacteria from oxidative stress. EMBO J. 33, 1896–1911 (2014).

14. T. W. Giessen, P. A. Silver, Widespread distribution of encapsulin nanocompartments reveals functional diversity. Nat. Microbiol. 2, 17029 (2017).

15. A. Tamura, Y. Fukutani, T. Takami, M. Fujii, Y. Nakaguchi, Y. Murakami, K. Noguchi, M. Yohda, M. Odaka, Packaging guest proteins into the encapsulin nanocompartment from Rhodococcus erythropolis N771. Biotechnol. Bioeng. 112, 13–20 (2015).

16. D. He, C. Piergentili, J. Ross, E. Tarrant, L. R. Tuck, C. L. Mackay, Z. McIver, K. J. Waldron, D. J. Clarke, J. Marles-Wright, Conservation of the structural and functional architecture of encapsulated ferritins in bacteria and archaea. Biochem. J. 476, 975–989 (2019).

17. R. Rahmanpour, T. D. H. Bugg, Assembly in vitro of Rhodococcus jostii RHA1 encapsulin and peroxidase DypB to form a nanocompartment. FEBS J. 280, 2097–2104 (2013).

18. J. Snijder, O. Kononova, I. M. Barbu, C. Uetrecht, W. F. Rurup, R. J. Burnley, M. S. T. Koay, J. J. L. M. Cornelissen, W. H. Roos, V. Barsegov, G. J. L. Wuite, A. J. R. Heck, Assembly and Mechanical Properties of the Cargo-Free and Cargo-Loaded Bacterial Nanocompartment Encapsulin. Biomacromolecules. 17, 2522–9 (2016).

19. M. P. Andreas, T. W. Giessen, Large-scale computational discovery and analysis of virus-derived microbial nanocompartments. Nat. Commun. 12, 4748 (2021).

20. M. C. Jenkins, S. Lutz, Encapsulin Nanocontainers as Versatile Scaffolds for the Development of Artificial Metabolons. ACS Synth. Biol. 10, 857–869 (2021).

21. Y. H. Lau, T. W. Giessen, W. J. Altenburg, P. A. Silver, Prokaryotic nanocompartments form synthetic organelles in a eukaryote. Nat. Commun. 9, 1311 (2018).

22. F. Sigmund, S. Pettinger, M. Kube, F. Schneider, M. Schifferer, S. Schneider, M. V. Efremova, J. Pujol-Martí, M. Aichler, A. Walch, T. Misgeld, H. Dietz, G.G. Westmeyer, Iron-Sequestering Nanocompartments as Multiplexed Electron Microscopy Gene Reporters. ACS Nano. 13, 8114–8123 (2019).

23. S. Recalcati, E. Gammella, P. Buratti, G. Cairo, Molecular regulation of cellular iron balance. IUBMB Life. 69, 389–398 (2017).

24. A. Tamura, Y. Fukutani, T. Takami, M. Fujii, Y. Nakaguchi, Y. Murakami, K. Noguchi, M. Yohda, M. Odaka, Packaging guest proteins into the encapsulin nanocompartment from Rhodococcus erythropolis N771. Biotechnol. Bioeng. 112, 13–20 (2015).

25. Y. Tang, A. Mu, Y. Zhang, S. Zhou, W. Wang, Y. Lai, X. Zhou, F. Liu, X. Yang, H. Gong, Q. Wang, Z. Rao, Cryo-EM structure of Mycobacterium smegmatis DyP-loaded encapsulin. Proc. Natl. Acad. Sci. 118, e2025658118 (2021).

26. M. Sutter, D. Boehringer, S. Gutmann, S. Günther, D. Prangishvili, M. J. Loessner, K. O. Stetter, E. Weber-Ban, N. Ban, Structural basis of enzyme encapsulation into a bacterial nanocompartment. Nat. Struct. Mol. Biol. 15, 939–947 (2008).

27. X. Xiong, C. Sun, F. S. Vago, T. Klose, J. Zhu, W. Jiang, Cryo-EM structure of heterologous protein complex loaded thermotoga maritima encapsulin capsid. Biomolecules. 10, 1–13 (2020).

28. R. M. Putri, C. Allende-Ballestero, D. Luque, R. Klem, K.-A. Rousou, A. Liu, C. H.-H. Traulsen, W. F. Rurup, M. S. T. Koay, J. R. Castón, J. J. L. M. Cornelissen, Structural Characterization of Native and Modified Encapsulins as Nanoplatforms for in Vitro Catalysis and Cellular Uptake. ACS Nano. 11, 12796–12804 (2017).

29. J. Ross, T. Lambert, C. Piergentili, D. He, K. J. Waldron, C. L. Mackay, J. Marles-Wright, D. J. Clarke, Mass spectrometry reveals the assembly pathway of encapsulated ferritins and highlights a dynamic ferroxidase interface. Chem. Commun. (Camb). 56, 3417–3420 (2020).

30. T. Nakane, E. Lindahl, J. Zivanov, W. J. J. H. Hagen, S. H. W. H. Scheres, D. Kimanius, B. B. O. Forsberg, T. Nakane, B. B. O. Forsberg, D. Kimanius, W. J. J. H. Hagen, E. Lindahl, S. H. W. H. Scheres, New tools for automated high-resolution cryo-EM structure determination in RELION-3. Elife. 7, 1–38 (2018).

31. R. J. Nichols, B. LaFrance, N. R. Phillips, D. R. Radford, L. M. Oltrogge, L. E. Valentin-Alvarado, A. J. Bischoff, E. Nogales, D. F. Savage, Discovery and characterization of a novel family of prokaryotic nanocompartments involved in sulfur metabolism. Elife. 10 (2021), doi:10.7554/eLife.59288.

32. B. LaFrance, C. Cassidy-Amstutz, R. J. Nichols, L. M. Oltrogge, E. Nogales, D. F. Savage, bioRxiv, in press, doi:10.1101/2021.04.26.441214.

33. E. M. Williams, S. M. Jung, J. L. Coffman, S. Lutz, Pore Engineering for Enhanced Mass Transport in Encapsulin Nanocompartments. ACS Synth. Biol. 7, 2514–2517 (2018).

34. A. Punjani, D. J. Fleet, 3D variability analysis: Resolving continuous flexibility and discrete heterogeneity from single particle cryo-EM. J. Struct. Biol. 213, 107702 (2021).

35. L. Adamson, N. Tasneem, M. P. Andreas, W. Close, T. N. Szyszka, E. Jenner Young, L. Chen Cheah, A. Norman, F. Sainsbury, T. W. Giessen, Y. Heng Lau, bioRxiv, in press, doi:10.1101/2021.01.27.428512.

36. C. Piergentili, J. Ross, D. He, K. J. Gallagher, W. A. Stanley, L. Adam, C. L. Mackay, A. Baslé, K. J. Waldron, D. J. Clarke, J. Marles-Wright, Dissecting the structural and functional roles of a putative metal entry site in encapsulated ferritins. J. Biol. Chem. (2020), doi:10.1074/jbc.RA120.014502.

37. S. V. Iverson, T. L. Haddock, J. Beal, D. M. Densmore, CIDAR MoClo: Improved MoClo Assembly Standard and New E. coli Part Library Enables Rapid Combinatorial Design for Synthetic and Traditional Biology. ACS Synth. Biol. 5, 151019092657002 (2015).

38. F. Studier, Protein production by auto-induction in high-density shaking cultures. Protein Expr. Purif. 41, 207–234 (2005).

39. J. Schindelin, I. Arganda-Carreras, E. Frise, V. Kaynig, M. Longair, T. Pietzsch, S. Preibisch, C. Rueden, S. Saalfeld, B. Schmid, J.-Y. Tinevez, D. J. White, V. Hartenstein, K. Eliceiri, P. Tomancak, A. Cardona, Fiji: an open-source platform for biological-image analysis. Nat. Methods. 9, 676–682 (2012).

40. S. Q. Zheng, E. Palovcak, J.-P. Armache, K. A. Verba, Y. Cheng, D. A. Agard, MotionCor2: anisotropic correction of beam-induced motion for improved cryo-electron microscopy. Nat. Methods. 14, 331–332 (2017).

41. A. Rohou, N. Grigorieff, CTFFIND4: Fast and accurate defocus estimation from electron micrographs. J. Struct. Biol. 192, 216–21 (2015).

42. A. Punjani, J. L. Rubinstein, D. J. Fleet, M. A. Brubaker, No Title. Nat. Methods. 14, 290–296 (2017).

43. T. D. Goddard, C. C. Huang, E. C. Meng, E. F. Pettersen, G. S. Couch, J. H. Morris, T. E. Ferrin, UCSF ChimeraX: Meeting modern challenges in visualization and analysis. Protein Sci. 27, 14–25 (2018).

44. L. A. Kelly, S. Mezulis, C. Yates, M. Wass, M. Sternberg, The Phyre2 web portal for protein modelling, prediction, and analysis. Nat. Protoc. 10, 845–858 (2015).

45. E. F. Pettersen, T. D. Goddard, C. C. Huang, E. C. Meng, G. S. Couch, T. I. Croll, J. H. Morris, T. E. Ferrin, UCSF ChimeraX: Structure visualization for researchers, educators, and developers. Protein Sci. 30, 70–82 (2021).

46. P. V. Afonine, B. K. Poon, R. J. Read, O. V. Sobolev, T. C. Terwilliger, A. Urzhumtsev, P. D. Adams, Real-space refinement in PHENIX for cryo-EM and crystallography. Acta Crystallogr. Sect. D Struct. Biol. 74, 531–544 (2018).

47. P. Emsley, B. Lohkamp, W. G. Scott, K. Cowtan, Features and development of Coot. Acta Crystallogr. D. Biol. Crystallogr. 66, 486–501 (2010).

48. V. B. Chen, W. B. Arendall, J. J. Headd, D. A. Keedy, R. M. Immormino, G. J. Kapral, L. W. Murray, J. S. Richardson, D. C. Richardson, MolProbity: all-atom structure validation for macromolecular crystallography. Acta Crystallogr. D. Biol. Crystallogr. 66, 12–21 (2010).

49. P. V. Afonine, B. P. Klaholz, N. W. Moriarty, B. K. Poon, O. V. Sobolev, T. C. Terwilliger, P. D. Adams, A. Urzhumtsev, IUCr, New tools for the analysis and validation of cryo-EM maps and atomic models. Acta Crystallogr. Sect. D Struct. Biol. 74, 814–840 (2018).

50. B. A. Barad, N. Echols, R. Y.-R. Wang, Y. Cheng, F. DiMaio, P. D. Adams, J. Fraser, EMRinger: side chain-directed model and map validation for 3D cryo-electron microscopy. Nat. Methods. 12, 943–6 (2015).

51. F. Sievers, D. G. Higgins, Clustal Omega, accurate alignment of very large numbers of sequences. Methods Mol. Biol. 1079, 105–16 (2014).

52. P. Gouet, X. Robert, E. Courcelle, ESPript/ENDscript: Extracting and rendering sequence and 3D information from atomic structures of proteins. Nucleic Acids Res. 31, 3320–3 (2003).

53. G. R. Masson, J. E. Burke, N. G. Ahn, G. S. Anand, C. Borchers, S. Brier, G. M. Bou-Assaf, J. R. Engen, S. W. Englander, J. Faber, R. Garlish, P. R. Griffin, M. L. Gross, M. Guttman, Y. Hamuro, A. J. R. Heck, D. Houde, R. E. Iacob, T. J. D. Jørgensen, I. A. Kaltashov, J. P. Klinman, L. Konermann, P. Man, L. Mayne, B. D. Pascal, D. Reichmann, M. Skehel, J. Snijder, T. S. Strutzenberg, E. S. Underbakke, C. Wagner, T. E. Wales, B. T. Walters, D. D. Weis, D. J. Wilson, P. L. Wintrode, Z. Zhang, J. Zheng, D. C. Schriemer, K. D. Rand, Recommendations for performing, interpreting and reporting hydrogen deuterium exchange mass spectrometry (HDX-MS) experiments. Nat. Methods. 16, 595–602 (2019).

54. M. Laguerre, M. Saux, J. P. Dubost, A. Carpy, in Pharmaceutical Sciences (John Wiley & Sons, Ltd, 1997), vol. 3, pp. 217–222.

