## Supplemental Data for "Pore dynamics and asymmetric cargo loading in an encapsulin nanocompartment"

**Supplementary Information: Pore dynamics and asymmetric cargo loading in an encapsulin nanocompartment revealed by Cryo-EM and HDX mass spectrometry**

**Supplementary Figures**

Figure S1: Purification of recombinant Empty-Enc and Loaded-Enc protein complexes

Figure S2: Negative stain TEM of purified Loaded-Enc and surrounding fractions

Figure S3: Supplementary cryo-EM data from the icosahedral reconstruction

Figure S4: Cryo-EM processing workflow

Figure S5: Supplementary cryo-EM data of Loaded-Enc

Figure S6: The four EncFtn environments within the icosahedral encapsulin nanocompartment viewed from the outside of the encapsulin nanocompartment

Figure S7: 3D variability of Loaded-Enc

Figure S8: Symmetry expansion EM maps

Figure S9: Model fit of the ‘open’ and ‘closed’ structures into the symmetry expansion maps

Figure S10: Comparison of the A-domain of ‘closed’ and ‘open’ pentamer structures

Figure S11: Sequence alignment of encapsulins and HK97

Figure S12: Electrostatic properties of the encapsulin nanocompartment pores

Figure S13: Sequence coverage of deuterium uptake of Empty-Enc and Loaded-Enc peptides

Figure S14: Deuterium fractional uptake of Loaded-Enc peptides

Figure S15: Differential HDX-MS analysis of Empty-Enc and Loaded-Enc

**Supplementary Tables**

### Table S1: Protein masses and assignments obtained by LC-MS

### Table S2: Cryo-EM data collection

Table S3: Cryo-EM data processing in Relion3.1

Table S4: Cryo-EM data processing in CryoSPARC

### Table S5: Model building and refinement of the ‘open’ and ‘closed’ pentamer conformations of the encapsulin nanocompartment

Table S6: Recorded uptake of deuterium for each peptide and timepoint observed in HDX-MS of Loaded-Enc

Table S7: Deuterium uptake for each peptide and timepoint observed in HDX-MS of Empty-Enc

### Table S8: gBlocks used in this study

Table S9: Protein constructs used in this study

### Table S10: Autoinduction media components

**Supplementary Movies**

Supplementary Movie 1: Views of 3DVA Mode 0

Supplementary Movie 2: Views of 3DVA Mode 1

Supplementary Movie 3: Views of 3DVA Mode 2

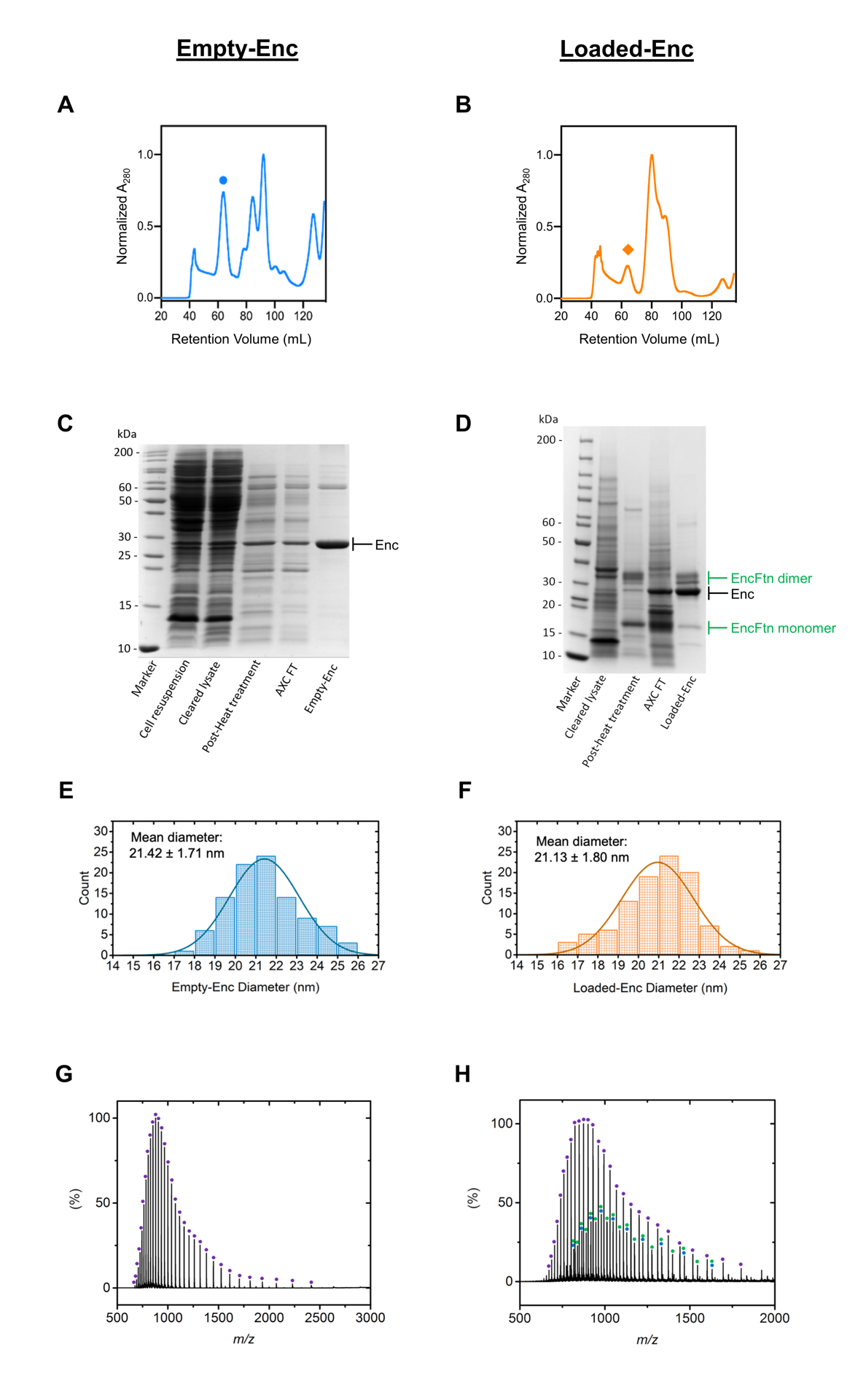

**Figure S1: Purification of recombinant *H. ochraceum* encapsulin protein complexes.**

**Top to Bottom: A/B:** Size exclusion chromatography traces of Empty-Enc (**A**) and Loaded-Enc (**B**) with normalized A_280_ values. Both Empty-Enc and Loaded-Enc elute with a retention volume of 65 mL (stressed with a blue circle and orange diamond respectively). Other peaks in the chromatogram represent smaller encapsulin oligomers and contaminants. The Loaded-Enc chromatogram has a peak at 80 mL with corresponds to excess unencapsulated EncFtn. **C/D**: SDS-PAGE gels (15% acrylamide stained with Coomassie blue stain) of the purification process of Empty-Enc (**C**) and Loaded-Enc (**D**). Anion exchange flow-through labelled as “AXC FT”, and concentrated and pooled SEC fractions labelled as “Empty-Enc” and “Loaded-Enc”. Encapsulin bands of approximately 30 kDa are labelled in black, and encapsulated ferritin bands (monomer at 15 kDa and dimer at 30 kDa) are labelled in green. The marker lane comprisesthe Fermentas PageRuler Unstained Protein Ladder. **E/F**: Histograms of the size distribution of Empty-Enc (**E**) and Loaded-Enc (**F**) from negative stain TEM. A Gaussian curve was fitted to the data by nonlinear least squares regression, showing that individual Empty-Enc nanocompartments have a mean diameter of 21.13 nm with a standard deviation of 1.80 nm, and Loaded-Enc have an average diameter of 21.42 nm with a standard deviation of 1.71 nm. **G/H**: Mass spectra of the proteins present in purified Empty-Enc (**G**) and Loaded-Enc (**H**). Empty-Enc displays one charge state distribution (highlighted with purple circles), which corresponds to the mass of the encapsulin monomer (28969.7 Da). Loaded-Enc shows three charge state distributions in agreement with the encapsulin nanocompartment protein (purple circles, 28813.2 Da), the EncFtn monomer (blue circles, 14667.4 Da), and the EncFtn dimer (green circles, 29334.5 Da).

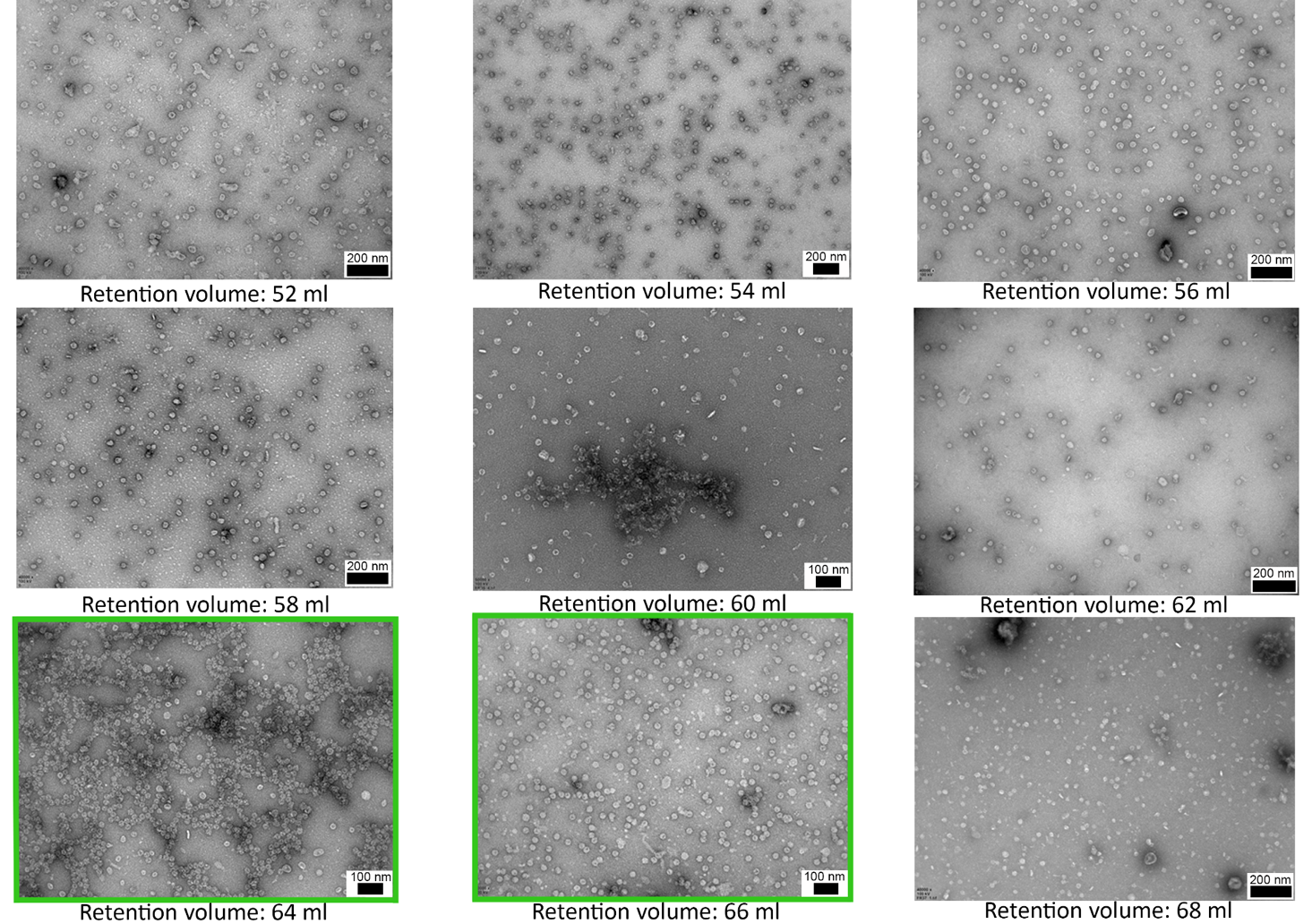

**Figure S2: Negative stain TEM of purified Loaded-Enc and surrounding size-exclusion chromatography fractions.** Indicative micrographs of fractions from the initial size exclusion chromatography purification step. Fractions are represented by their centred retention volume (mL). Fractions included in the final pool, for further analysis are highlighted in green. Scale bars are represented on each image individually.

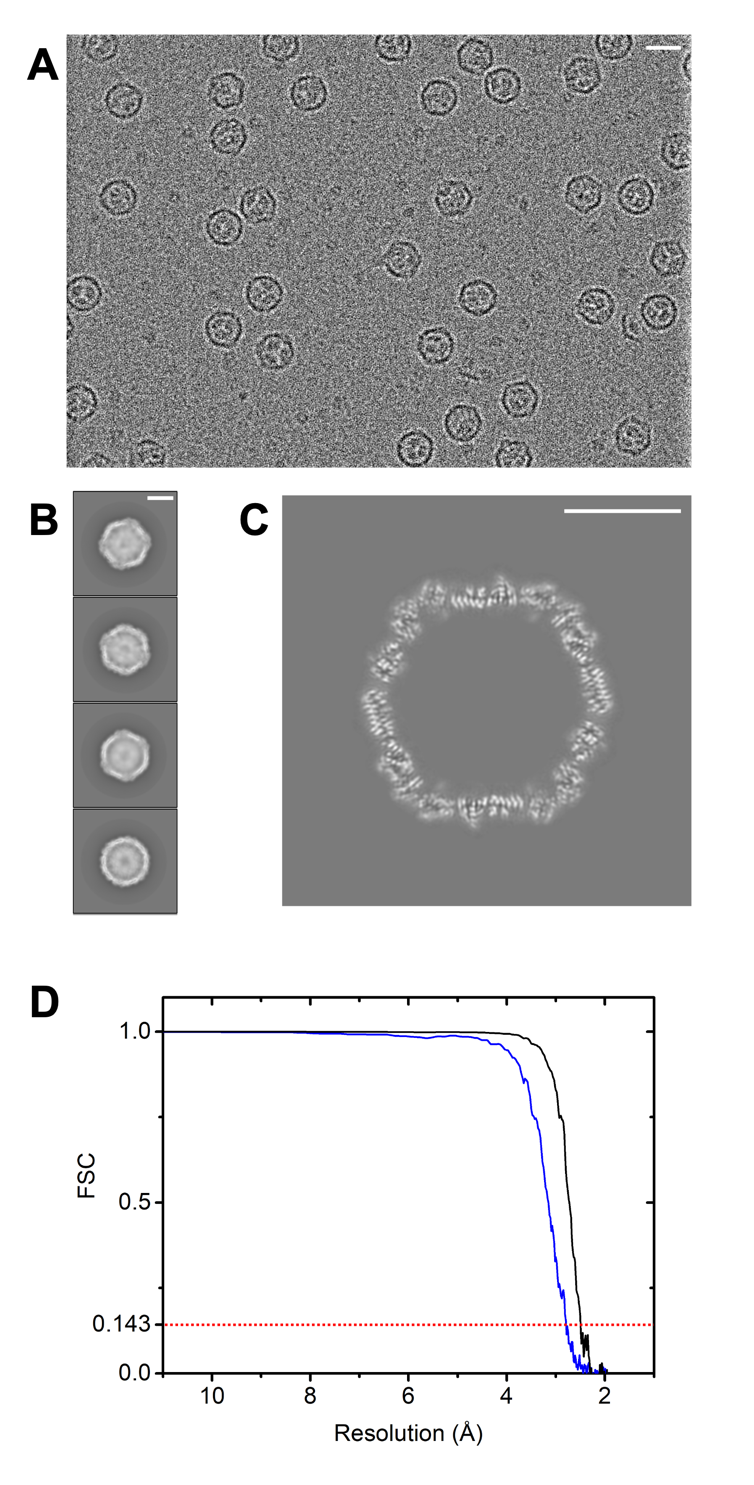

**Figure S3: Representative cryo-EM micrograph and additional data from the icosahedral reconstruction of the *H. ochraceum* encapsulin complex.**

**A**: Representative cryo-EM micrograph of Loaded-Enc. **B**: 2D classes of Loaded-Enc **C**: Central slice of Loaded-Enc from icosahedral processing. A white scale bar representing 10 nm is shown in the upper right corner of **A**, **B** and **C**. **D**: Gold standard FSC curve of the masked (black) and unmasked (blue) icosahedral reconstruction half maps. The FSC 0.143 threshold is highlighted with a dashed red line.

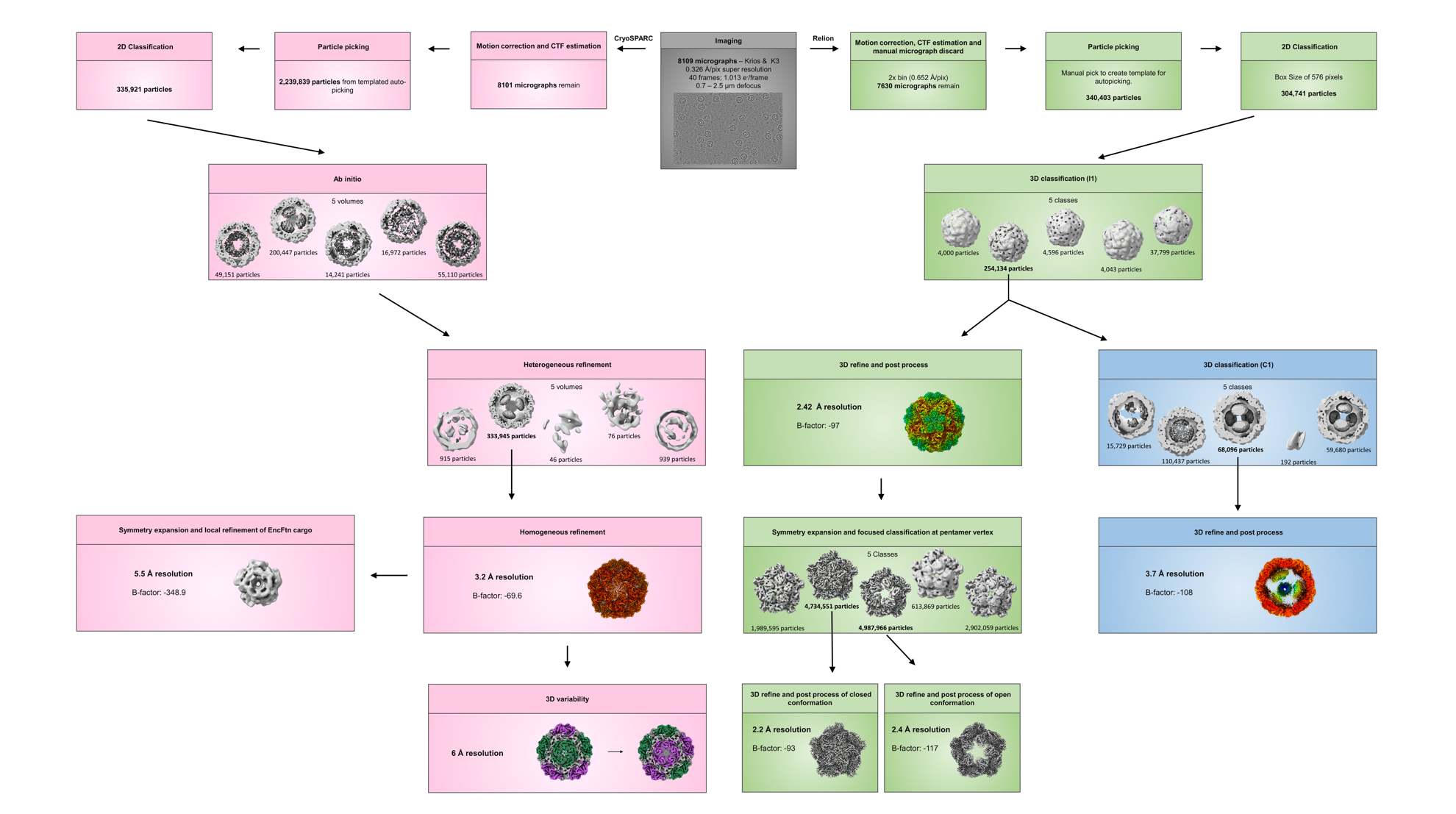

**Figure S4: Cryo-EM processing workflow.**

The processing pipeline within Relion3.1 and CryoSPARC used to obtain the single particle reconstructions of Loaded-Enc. The green boxes show the processing workflow for the initial processing steps and I1 reconstruction. The blue boxes show the fork to C1 processing to gain insight into the EncFtn loading inside Enc. Pink boxes show the CryoSPARC pipeline used for 3D variability analysis, and for symmetry expansion with local refinement to improve the resolution of the EncFtn cargo.

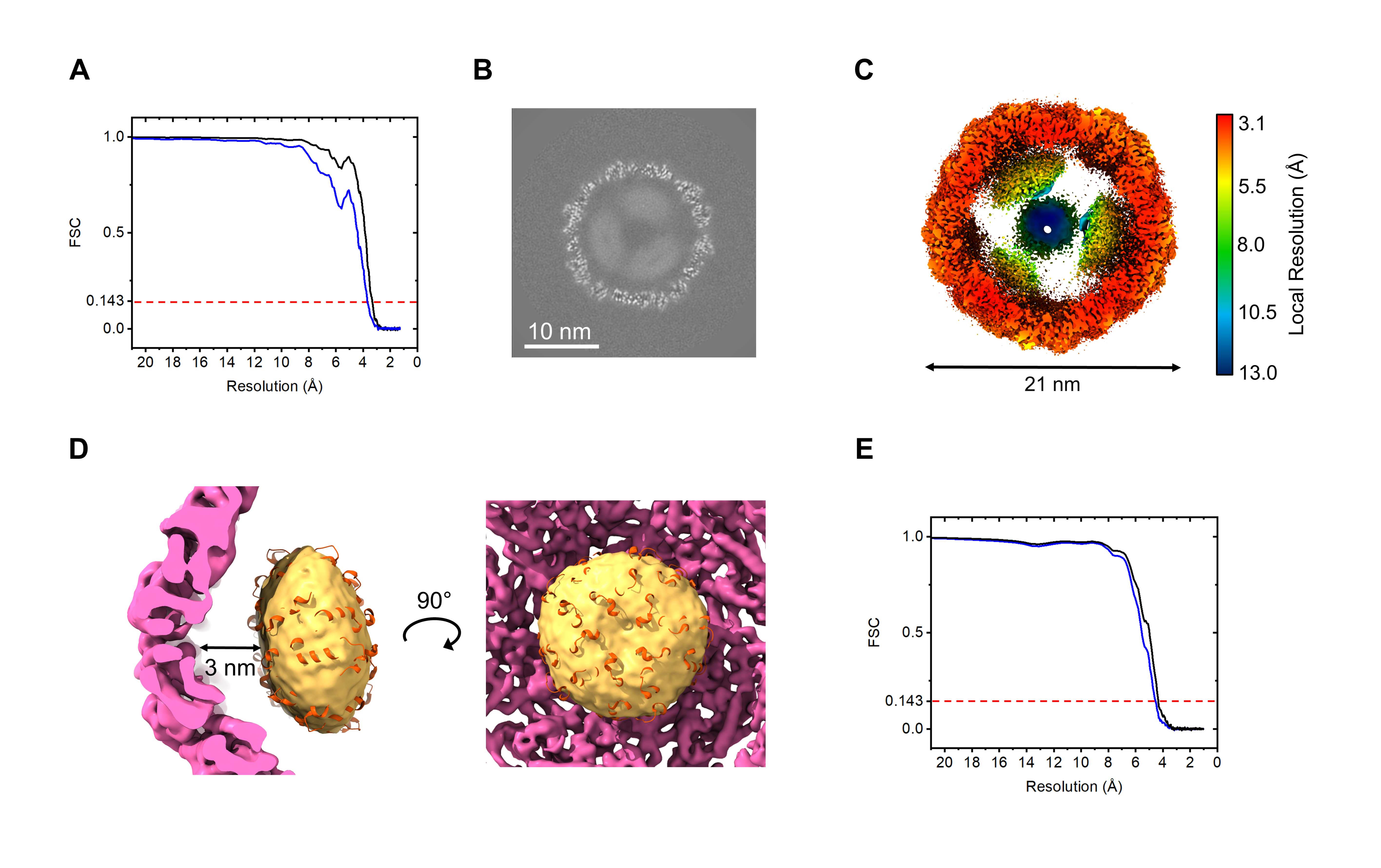

**Figure S5: Supplementary cryo-EM data of Loaded-Enc C1 reconstruction.**

**A**: Gold standard FSC curve of the masked (black, spherical mask) and unmasked (blue) icosahedral reconstruction half maps. The FSC 0.143 threshold is highlighted with a dashed red line. **B**: Central slice of Loaded-Enc from C1 reconstruction after 3D refinement. A white scale bar representing 10 nm is shown in the lower left corner. **C:** Slice through the locally filtered and sharped EM map of Loaded-Enc is shown colored by local resolution. The interior of the nanocompartment holds four EncFtn which are of significantly lower resolution than the shell. The color key for the local resolution is shown on the right side of the figure. **D:** The crystal structure of the EncFtn from *H. ochraceum* (orange, PDB 5N5F) docked into the interior density (yellow) of the C1 reconstruction. A 3 nm gap is observed between the Enc shell (pink) and the EncFtn (yellow and orange). **E**: The GS-FSC for the EncFtn cargo reconstruction produced by CryoSPARC. Non-masked shown in blue, spherical mask in black. 0.143 cut-off in red.

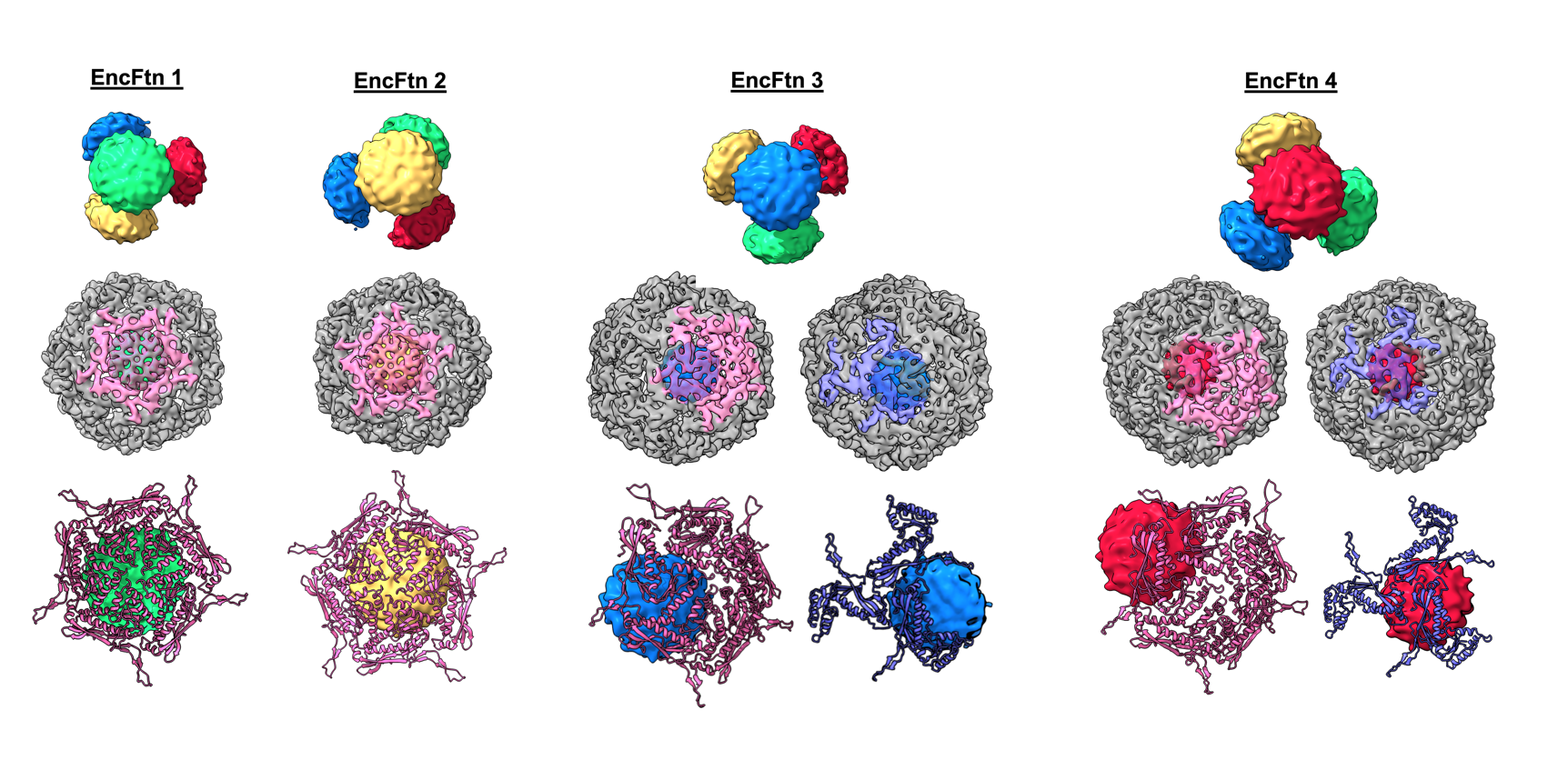

**Figure S6: Visualization of the distinct EncFtn environments within the icosahedral encapsulin nanocompartment.**

**Top panels:** Each EncFtn within the encapsulin nanocompartment has been individually colored (as in **Figure 4 C**) and numbered. Four orientations of EncFtn tetrahedral are shown with a different EncFtn in the foreground. **Middle panels:** Each EncFtn of the Loaded-Enc complex as viewed from outside of the encapsulin nanocompartment. EncFtn complexes are in the same orientation as the top panels. A pentamer of the encapsulin nanocompartment has been colored pink to allow direct correlation between the EncFtn location and the 5-fold pore of encapsulin. EncFtn 3 and EncFtn 4 do not align with the 5-fold pore and so the encapsulin 3-fold pore has also been shown and colored purple. **Bottom panels:** The relationship between each EncFtn and the pores of encapsulin. The encapsulin 5-fold pore is shown by pink cartoons, the encapsulin 3-fold pore by purple cartoons and the EncFtn are colored as in the top panels. EncFtn 1 and EncFtn 2 are in broadly equivalent environments which are aligned with the 5-fold pore of the encapsulin nanocompartment.

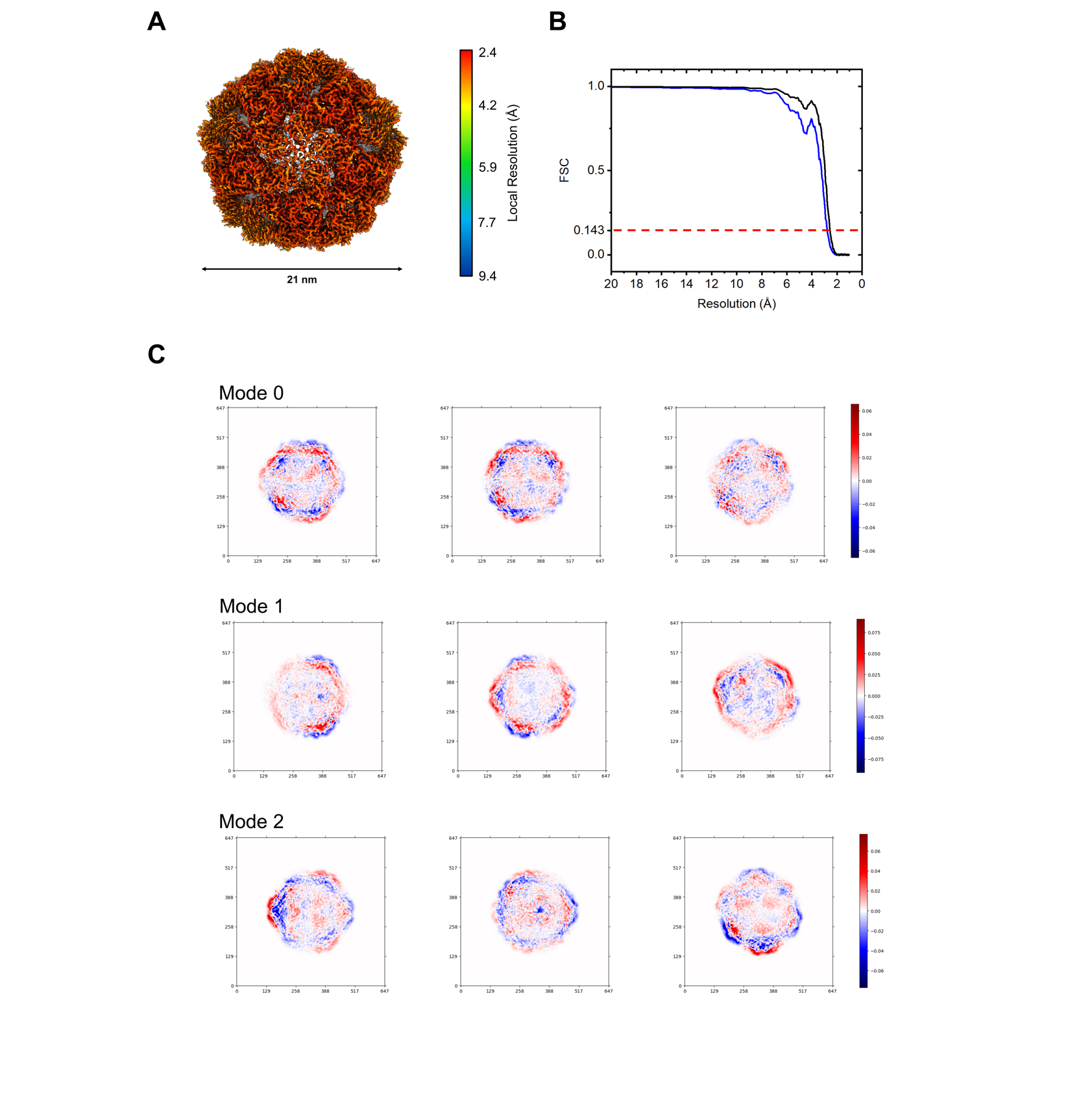

**Figure S7: 3D variability analysis of Loaded-Enc. A:** Density map of the consensus C1 reconstruction of Loaded-Enc calculated using CryoSPARC is shown colored by local resolution. The color key for the local resolution is shown on the right side of the panel. **B**: Gold standard FSC curve of the masked (spherical mask, black) and unmasked (blue) CryoSPARC reconstruction half maps. The GS-FSC 0.143 threshold is highlighted with a dashed red line. **C**: 2D projections of orthogonal slices of the three variability modes calculated.

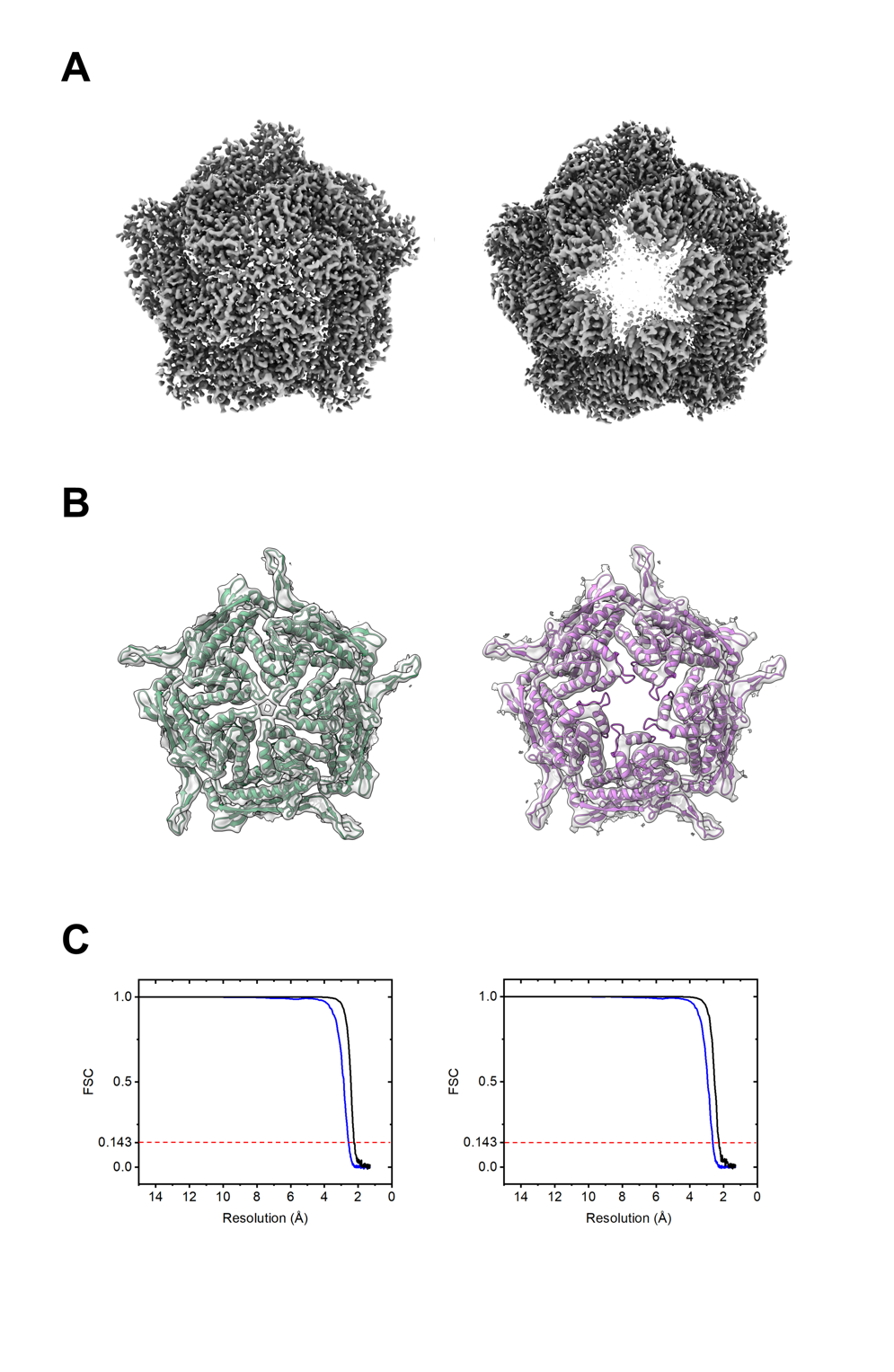

**Figure S8: Electronic potential maps and models from masked 3D-classification and refinement centered on the five-fold symmetry axis of the symmetry expanded icosahedral reconstruction.**

**A:** Cryo-EM maps of the ‘closed’ (left) and ‘open’ (right) pentamer conformations from symmetry expansion of the icosahedral five-fold axis. **B**: The Gaussian smoothed cryo-EM maps with docked models of the ‘closed’ (green, left) and ‘open’ (purple, right) conformations. Smoothed EM maps allow easy visualization of the docked secondary structure. **C**: GS-FSC for closed (left panel) and open (right panel) conformations of the encapsulin nanocompartment 5-fold pore.The 0.143 GS cut-off has been highlighted with a dashed red line. Curve for masked maps shown in black, and unmasked in blue.

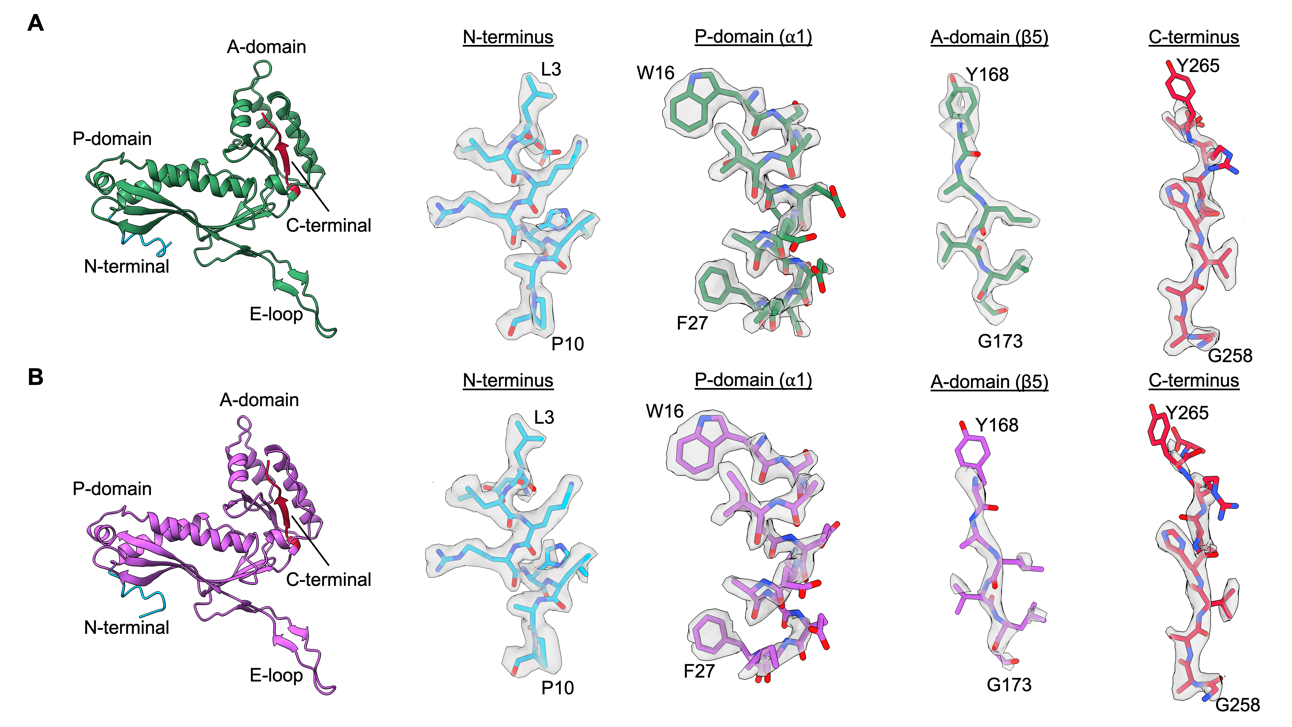

**Figure S9: Model fit of the ‘open’ and ‘closed’ atomic models produced from open and closed maps.**

The atomic models of the monomer subunit of the ‘closed’ (green, **A**) and ‘open’ (purple, **B**) pentamer conformations. Representative maps are shown across the monomer chain illustrating the fit of the atomic model and side chains into the cryo-EM map.

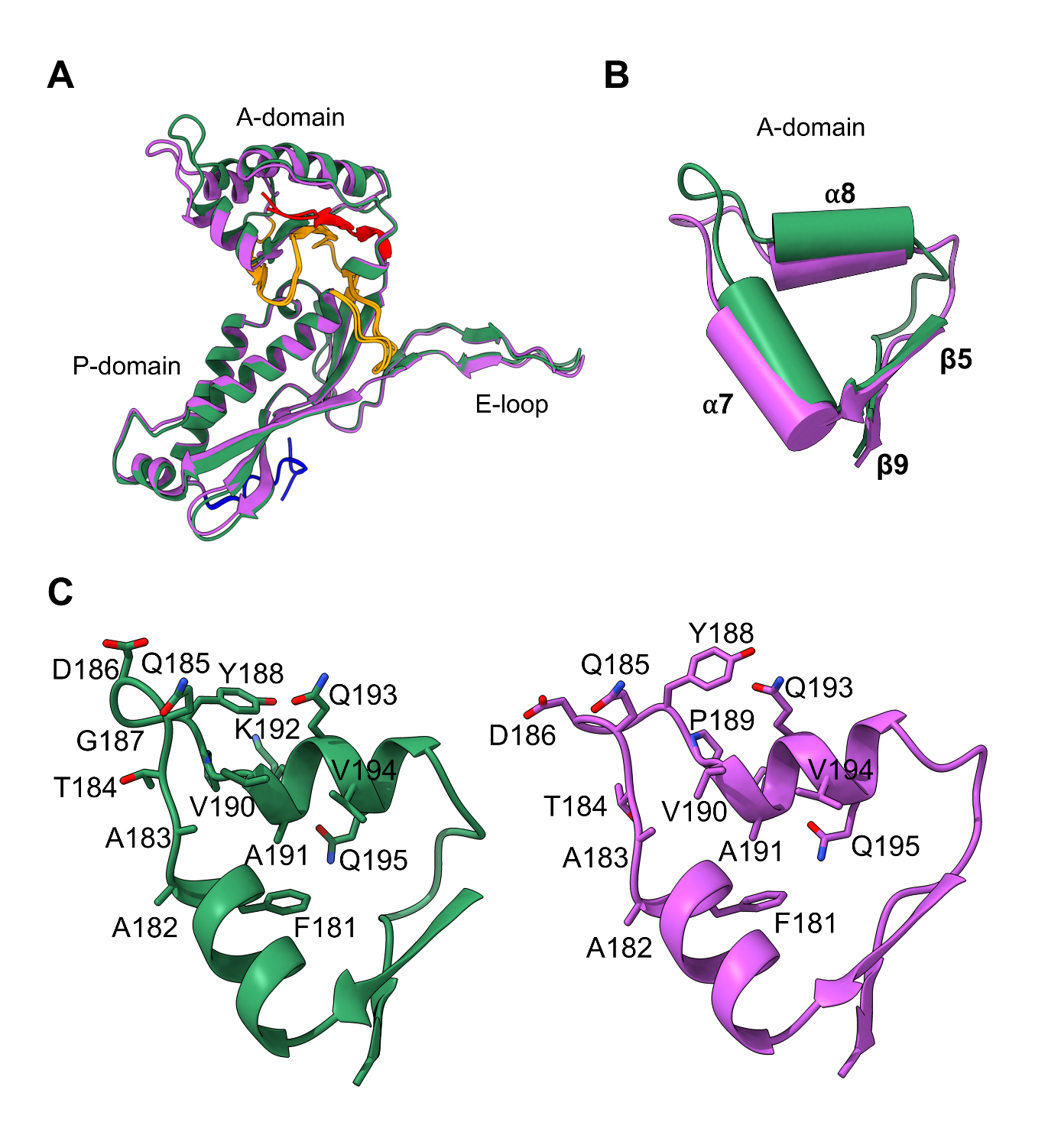

**Figure S10: Comparison of the A-domain of ‘closed’ and ‘open’ pentamer structures.**

**A**: Overlay of the monomers from the ‘closed’ (green) and ‘open’ (purple) pentamer conformations displaying shifted A-domains. The hinge regions between the P- and A-domains are colored orange, and the N- and C-termini are colored orange, with N- and C-termini coloured in blue and red respectively. **B**: Overlay of the A-domain of the ‘closed’ (green) and ‘open’ (purple) conformations highlighting the change in alpha helices 7 and 8. **C**: The residues present at the 5-fold pore and in alpha helices 7 and 8.

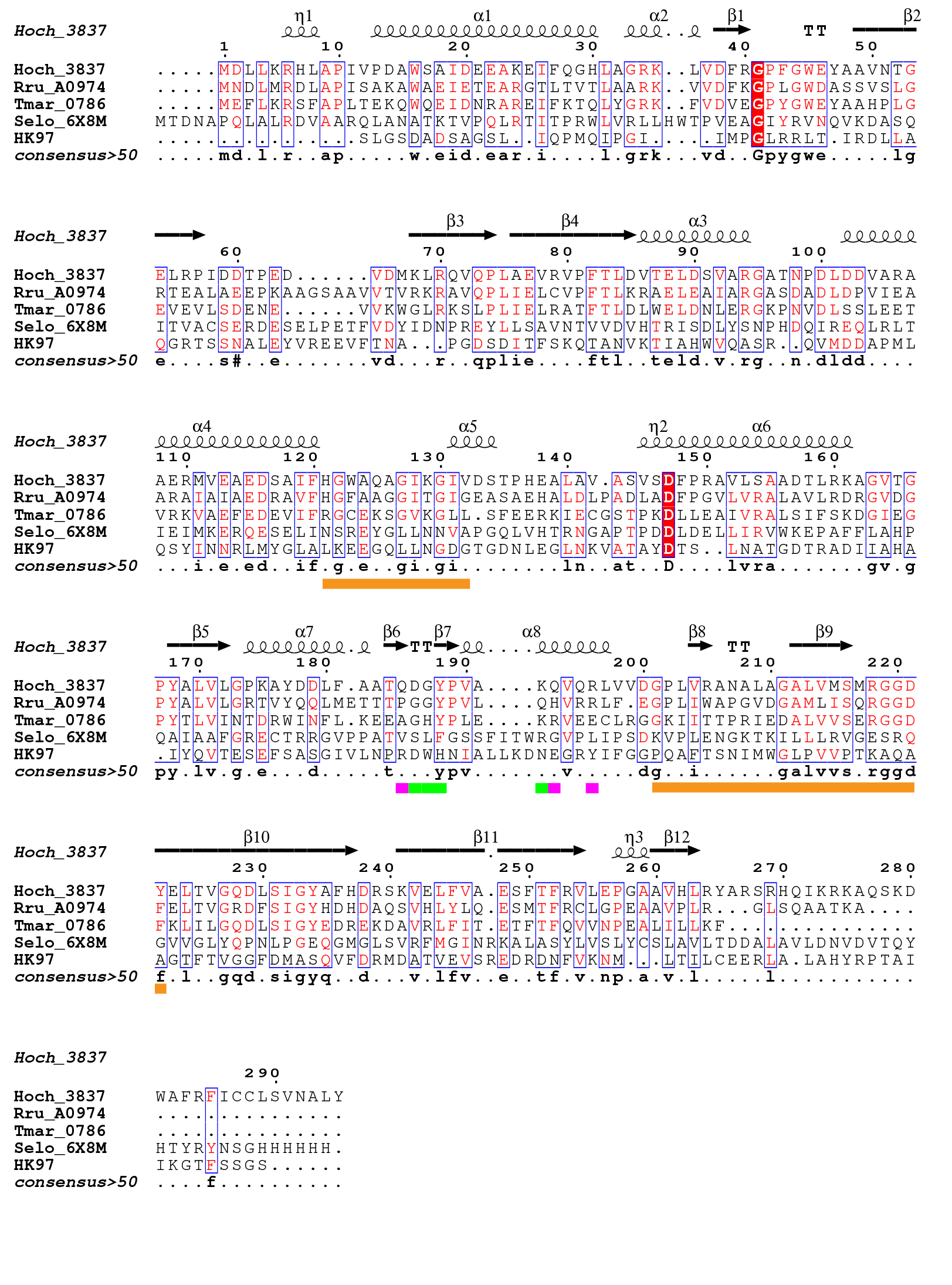

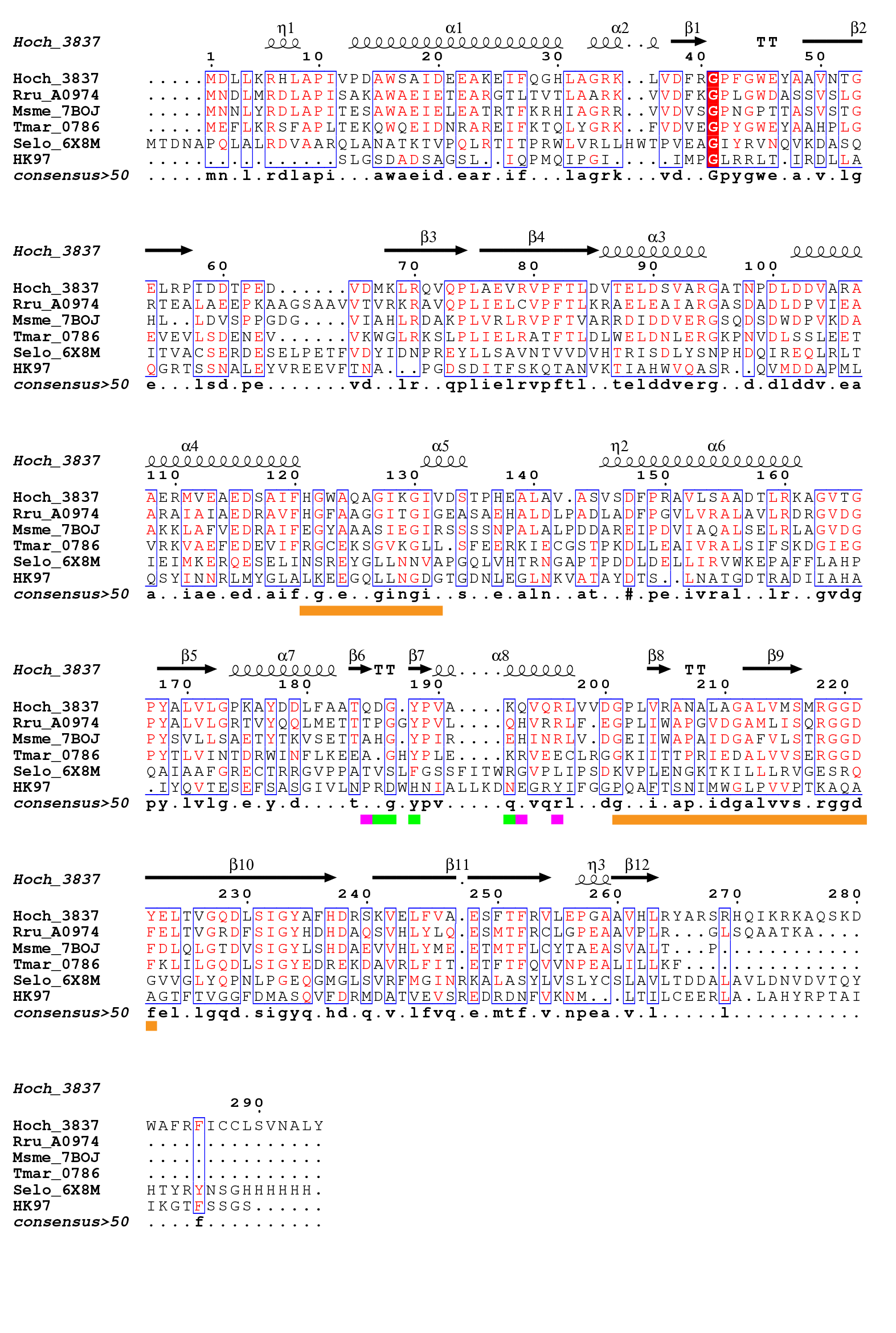

**Figure S11: Sequence alignment of encapsulins and HK97.**

Protein sequence alignment of encapsins from the *Haliangium ochraceum* (Hoch_3837), *Rhodospirillum rubrum* (Rru_A0974), *Mycobacterium smegmatis* (Msme_7BOJ), *Thermotoga maritima* (Tmar_0786) and *Synechococcus* *elongatus* (Selo_6X8M). Encapsulins share the HK97-fold from the HK97 bacteriophage capsid which is also shown in the sequence alignment (HK97). The residues in the 5-fold pore in the ‘closed’ pentamer conformation are underlined in purple. The additional residues which become exposed and form the 5-fold pore in the ‘open’ conformation are underlined in green. Residues in the hinge regions between the A- and P-domains are underlined in orange. Protein sequences were sourced from uniprot and the alignment was performed with Clustal Omega, and then formatted using ESPript.

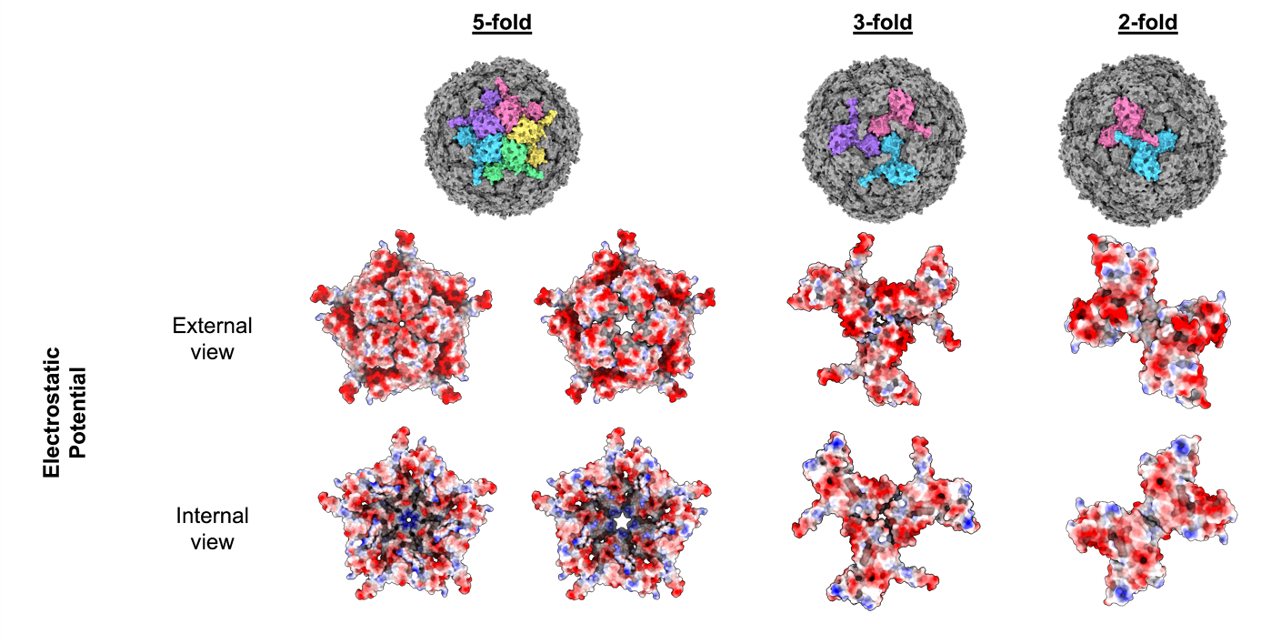

**Figure S12: Electrostatic properties of the encapsulin nanocompartment pores.**
From top to bottom: The top row shows a T=1 encapsulin in three orientations to show the 5-fold, 3-fold and 2-fold symmetry axes. Monomers have been colored individually (pink, yellow, green, blue, and purple) to highlight the symmetry axes. The second and third row show the 5-fold (closed and open conformations), 3-fold and 2-fold Loaded-Enc symmetry axes colored with electrostatic surfaces (positive charges shown in red and negative in blue).

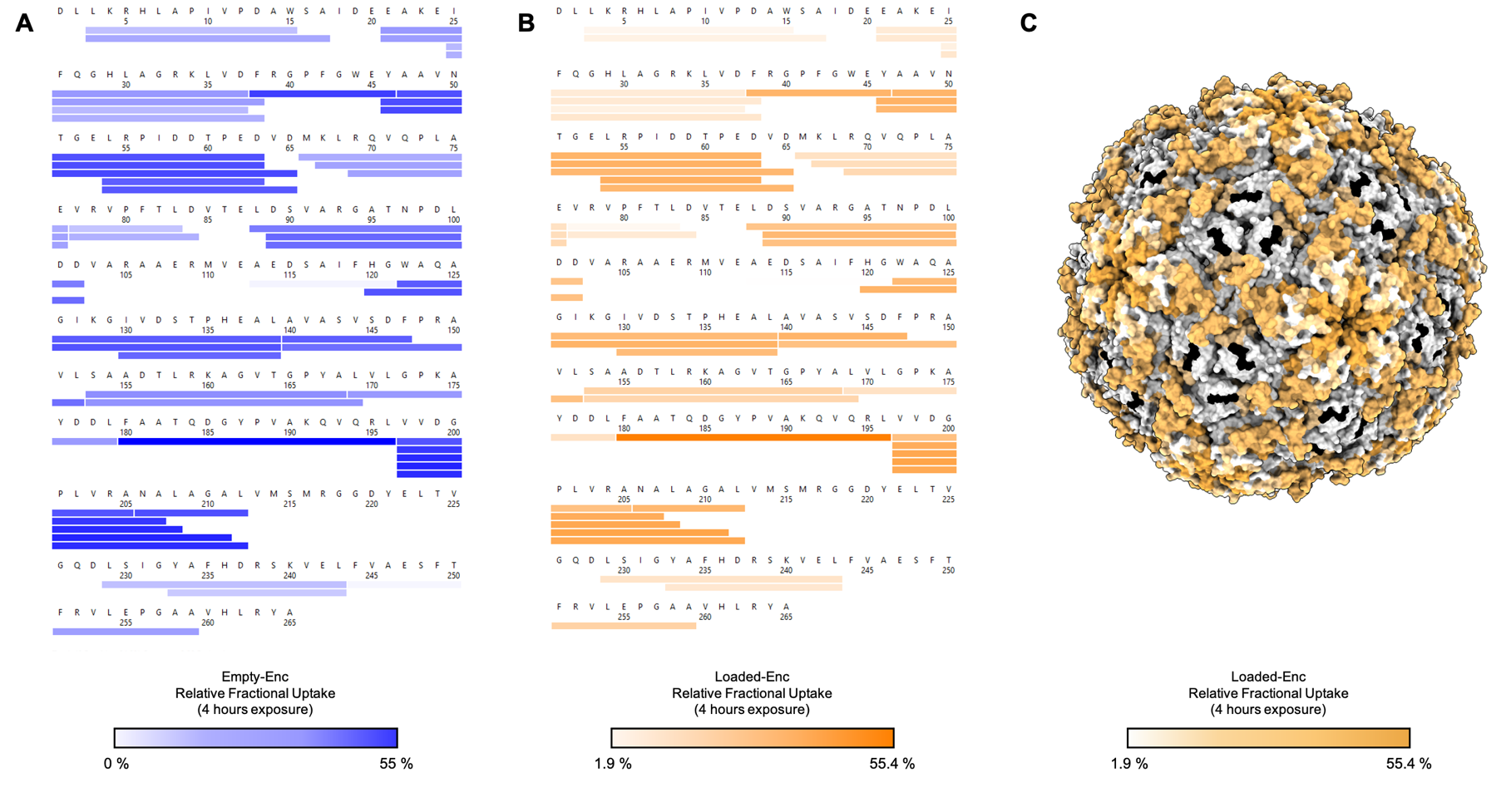

**Figure S13: Sequence coverage and deuterium uptake after HDX-MS analysis of Empty-Enc and Loaded-Enc peptides.**

HDX coverage maps showing the relative fractional deuterium uptake for both Empty-Enc (**A**) and Loaded-Enc (**B**) after 4 hours of deuterium labelling. There are 40 observed peptides common to both states, providing 84 % protein sequence coverage with 2.35 redundancy. Color keys are shown under **A** and **B**. (**C**) The relative fractional uptake of Loaded-Enc displayed on the encapsulin nanocompartment. Areas colored black are representative of no sequence coverage

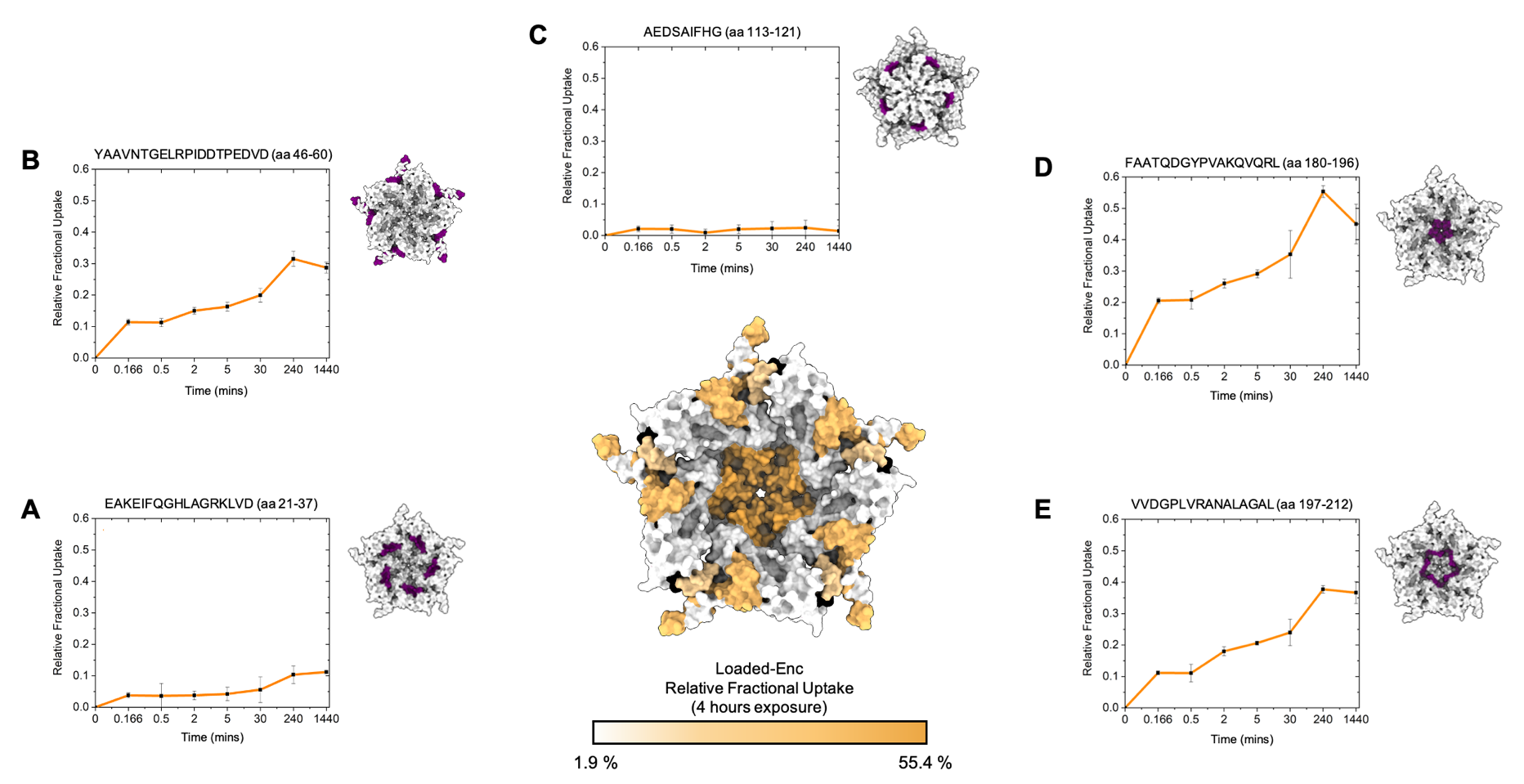

**Figure S14: Deuterium fractional uptake of Loaded-Enc peptides.**

HDX-MS of the encapsulin 5-fold pore highlighting the amount of deuterium incorporation after 4 hours of exposure, colored according to uptake (a color key is shown at the bottom of the central figure). (**A**) to (**E**), Uptake plots for individual peptides showing the relative deuterium uptake over time. An encapsulin pentamer is shown with the corresponding peptide sequence colored purple. (**A**) residues 21-37, whose proximity is close to the proposed localization sequence for EncFtn (11.2 ± 0.2 % uptake at 4 hours). (**B**) residues 46-60, which overlays with the proposed 2-fold symmetry pore (31.7 ± 2.2 % uptake at 4 hours), (**C**) residues 113-121, a highly protected exterior peptide (0.2 ± 0 % at 4 hours), (**D**) residues 180-196 the potential 5-fold symmetry pore (55.4 ± 2.2 % at 4 hours), (**E**) interior residue 197-212 in close proximity to the potential 5-fold symmetry pore site (37.7 ± 1.2 % at 4 hours).

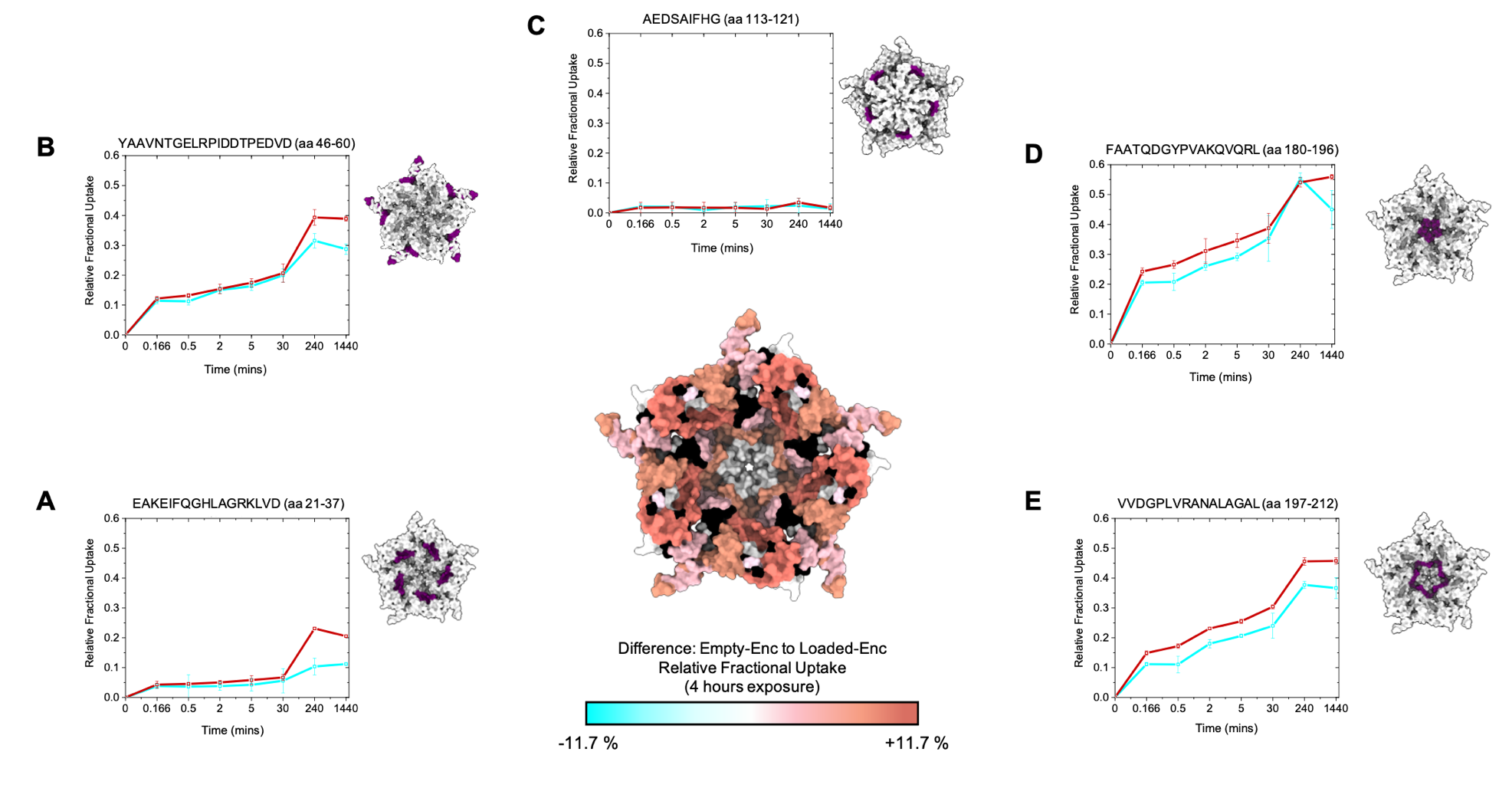

**Figure S15: Differential HDX-MS analysis of Empty-Enc and Loaded-Enc.**

The amount of deuterium incorporation over time is shown for five representative peptides (**A** - **E**) for both Empty-Enc (r lines) and Loaded-Enc (cyan lines). An encapsulin pentamer is shown next to each uptake chart with the corresponding peptide colored in purple. The central figure shows the difference in exchange is displayed on the structure of the 5-fold pore of encapsulin, with a color key below and areas of no sequence coverage colored black. (**A**) Location of residues 21-37 difference of +11.7 ± 1.5 %, (**B**) residues 46-65 with a difference of +7.6 ± 1.8 % (**C**) Location of residues 113-121 with a difference of 0.1 ± 0 %, (**D**) residues 180-196 with a difference of 0 ± 0.3 %, (**E**) interior residue 197-212 with a difference of 7.9 ± 1.7 %.

### Table S1: Protein masses and assignments obtained by LC-MS

| **Sample** | **Protein in Sample** | **Observed Average Mass (Da)** | **Assignment** | **Theoretical Average Mass (Da)** |
| --- | --- | --- | --- | --- |
| Empty-Enc | Encapsulin from *H. ochraceum* | 28969.67 ± 0.23 | Hoch-Enc monomer | 28968.84* |
| Loaded-Enc | Encapsulin from *H. ochraceum* | 28813.24 ± 0.89 | Hoch-Enc monomer | 28812.66* |
|  | EncFtn from *H. ochraceum* | 14667.42 ± 0.22 | Hoch-EncFtn monomer without Met | 14668.20 |
|  |  | 29334.50 ± 0.64 | Hoch-EncFtn dimer without Met | 29336.40 |

*The starting methionine residue is not always retained, and this has been indicated in “Assignment”. Errors generated by MassLynx v4.1.*

*** The difference in theoretical mass for the two *H. ochraceum* encapsulin proteins is due to an additional C-terminal arginine residue in the Empty-Enc construct which is a cloning artefact from the MoClo kit used.

### Table S2: Cryo-EM data collection

| **Data Collection** | |
| --- | --- |
| Microscope | FEI Titan Krios (300 kV) |
| Detector | Gatan K3 |
| Acquisition Mode | Super resolution |
| Pixel Size | 0.326 Å/pix (super-resolution)  0.652 Å/pix (physical) |
| Total dose | 40.509 e^-^/Å^2^ |
| Fractional dose | 40 frames during a 1 s exposure |
| Defocus range | 0.7 – 2.5 μm (0.3 μm steps) |
| Micrograph movies recorded | 8109 |
| EMPIAR entry | EMPIAR-1218 |

### Table S3: Cryo-EM data processing in Relion3.1

|  | **Loaded-Enc Icosahedral Reconstruction** | **Loaded-Enc**  **C1 Reconstruction** |
| --- | --- | --- |
| **Reconstruction in Relion** | | |
| Alignment Software | MotionCor2 | |
| Dose weighting | yes | |
| CTF fitting software | CTFFIND4 | |
| Correction | full | |
| Particle picking method/software | Relion – templated from manual picked 2D classes | |
| Particles picked | 340,403 | |
| Particles in 2D classes | 304,741 | |
| Particles used in final 3D reconstruction | 254,134 | 68,096 |
| Alignment software | Relion 3.1 | |
| Reconstruction software | Relion 3.1 | |
| Box size | 576 pixels (376 Å) | 480 pixels  (313 Å) |
| Voxel size (Å) | 0.652 | 0.652 |
| Symmetry | I1 | C1 |
| Map resolution (GS-FSC 0.143) (Å) | 2.5 | 3.7 |
| Sharpening B-factor (Å^2^) | -97 | -108 |
| EMDB ID | EMD-12853 | EMD-12873 |

**Table S4: Cryo-EM data processing in CryoSPARC**.

| **Reconstruction in CryoSPARC** | |
| --- | --- |
| Alignment method | Patch motion correction |
| CTF fitting method | Patch CTF |
| Particle picking method | Blob picker for initial picking  Template picking for subsequent picking |
| Particles picked | 472,247 |
| Particles in 2D classes | 335,921 |
| Particles after heterogeneous and homogeneous refinement | 333,945 |
| Particles used in 3D variability of encapsulin nanocompartment | 335,921 |
| Particles used in symmetry expand for EncFtn refinement | 89,398  (volume 2 from homogenous refine) |
| Particles used in local refinement of EncFtn | 357,592 (output from symmetry expand) |
| Box size | 648 pixels |
| Voxel size (Å) | 0.652 |
| Symmetry | C1 |
| **Loaded-Enc nanocompartment reconstruction** | |
| Map resolution (GS-FSC 0.143) (Å) | 3.2 |
| Sharpening B-factor (Å^2^) | -69.6 |
| EMDB ID | 13603 |
| **Loaded-Enc EncFtn cargo reconstruction** | |
| Map resolution (GS-FSC 0.143) (Å) | 5.5 |
| Sharpening B-factor (Å2) | -348.9 |
| EMDB ID | 13608 |

### Table S5: Model building and refinement of the ‘open’ and ‘closed’ pentamer conformations of the encapsulin nanocompartment.

| **Symmetry expansion of I1 dataset** | | |
| --- | --- | --- |
|  | **Closed Enc pentamer** | **Open Enc pentamer** |
| **Reconstruction** | | |
| Number of asymmetric units in final model | 4,734,551 | 4,987,966 |
| Map resolution (GS-FSC 0.143) (Å) | 2.2 | 2.4 |
| Map Sharpening B-factor (Å^2^) | -93 | -117 |
| EMDB ID | EMD-12859 | EMD-12864 |
| **Coordinate refinement** | | |
| Software | Phenix | Phenix |
| Refinement algorithm | Real space | Real space |
| Resolution cutoff (Å) | 2.2 | 2.4 |
| FSC_model-vs-map_= 0.5 (Å) | 2.3 | 2.5 |
| **Model** | | |
| Number of amino acid residues | 1375 | 1375 |
| Bond length outliers | 0 | 0 |
| Bond angle outliers | 0 | 0 |
| RMS deviations | | |
| Bonds (Å) | 0.006 | 0.004 |
| Angles (˚) | 0.674 | 0.638 |
| **Validation** | | |
| Molprobity score | 1.41 | 1.64 |
| Clash score | 3.13 | 6.34 |
| Rotamer outliers (%) | 2.2 | 0 |
| C_b_ outliers (%) | 0 | 0 |
| CaBLAM outliers (%) | 1.1 | 1.5 |
| Ramachandaran (%)  Favoured  Outlier | 97.79  2.21 | 97.05  2.95 |
| Model vs Data CC (mask) | 0.87 | 0.82 |
| EM Ringer score | 5.66 | 4.88 |
| PDB ID | 7OE2 | 7OEU |

**Table S6: Recorded uptake of deuterium for each peptide and timepoint observed in HDX-MS of Loaded-Enc.** For each peptic peptide observed in the HDX-MS analysis, the deuterium uptake (in Da) and standard deviation (SD) of triplicate data is shown for each timepoint (10 s, 30 s, 2 mins, 5 mins, 4 hours and 24 hours) for each peptide. The number of exchangeable backbone amide hydrogens is also stated (exchangers).

|  |  | **Loaded-Enc deuterium uptake (Da)** | | | | | | | | | | | | | | | |
| --- | --- | --- | --- | --- | --- | --- | --- | --- | --- | --- | --- | --- | --- | --- | --- | --- | --- |
| **Sequence** | **Exchangers** | **Start** | **End** | **10s** | **10s SD** | **30s** | **30s SD** | **2 mins** | **2 mins SD** | **5 mins** | **5 mins SD** | **30 mins** | **30 mins SD** | **4 hrs** | **4 hrs SD** | **24 hrs** | **24 hrs SD** |
| LKRHLAPIVPDAW | 10 | 3 | 15 | 0.34 | 0.08 | 0.31 | 0.10 | 0.36 | 0.29 | 0.36 | 0.15 | 0.38 | 0.08 | 0.55 | 0.08 | NaN | NaN |
| LKRHLAPIVPDAWSA | 12 | 3 | 17 | 0.45 | 0.06 | 0.39 | 0.08 | 0.36 | 0.09 | 0.41 | 0.06 | 0.56 | 0.11 | 0.89 | 0.05 | NaN | NaN |
| EAKEIFQGHLAGRKLVD | 16 | 21 | 37 | 0.62 | 0.06 | 0.55 | 0.10 | 0.59 | 0.10 | 0.65 | 0.08 | 0.90 | 0.14 | 1.66 | 0.06 | 1.42 | 0.19 |
| EAKEIFQGHLAGRKLVDF | 17 | 21 | 38 | 0.62 | 0.03 | 0.56 | 0.06 | 0.62 | 0.06 | 0.79 | 0.05 | 1.08 | 0.22 | 1.90 | 0.03 | 1.52 | 0.16 |
| IFQGHLAGRKLVD | 12 | 25 | 37 | 0.34 | 0.08 | 0.32 | 0.08 | 0.31 | 0.11 | 0.39 | 0.09 | 0.43 | 0.09 | 0.95 | 0.13 | 0.69 | 0.13 |
| IFQGHLAGRKLVDF | 13 | 25 | 38 | 0.40 | 0.05 | 0.38 | 0.05 | 0.44 | 0.06 | 0.61 | 0.08 | 0.79 | 0.20 | 1.58 | 0.10 | NaN | NaN |
| FRGPFGWEY | 7 | 38 | 46 | 0.82 | 0.06 | 0.70 | 0.28 | 0.93 | 0.10 | 0.97 | 0.15 | 1.45 | 0.29 | 2.30 | 0.07 | 1.89 | 0.03 |
| YAAVNTGELRPIDDTPED | 15 | 46 | 63 | 1.83 | 0.13 | 1.72 | 0.44 | 2.32 | 0.18 | 2.54 | 0.10 | 3.11 | 0.57 | 5.13 | 1.70 | 4.05 | 0.34 |
| YAAVNTGELRPIDDTPEDVD | 17 | 46 | 65 | 1.92 | 0.11 | 1.89 | 0.45 | 2.53 | 0.15 | 2.75 | 0.15 | 3.37 | 0.66 | 5.40 | 0.20 | 4.52 | 0.50 |
| AAVNTGELRPIDDTPED | 14 | 47 | 63 | 1.81 | 0.05 | 1.66 | 0.45 | 2.41 | 0.13 | 2.54 | 0.14 | 3.04 | 0.47 | 4.63 | 0.18 | 3.68 | 0.27 |
| LRPIDDTPED | 7 | 54 | 63 | 0.97 | 0.08 | 0.97 | 0.29 | 1.20 | 0.15 | 1.29 | 0.05 | 1.70 | 0.29 | 2.48 | 0.11 | 1.94 | 0.08 |
| LRPIDDTPEDVD | 9 | 54 | 65 | 1.13 | 0.12 | 0.99 | 0.31 | 1.35 | 0.18 | 1.53 | 0.14 | 1.83 | 0.28 | 2.87 | 0.12 | 2.59 | 0.24 |
| MKLRQVQPLAE | 9 | 66 | 76 | 0.82 | 0.12 | 0.73 | 0.23 | 0.67 | 0.15 | 0.75 | 0.23 | 1.01 | 0.14 | 1.29 | 0.08 | 1.17 | 0.23 |
| KLRQVQPLAE | 8 | 67 | 76 | 0.88 | 0.09 | 0.78 | 0.20 | 0.81 | 0.12 | 0.87 | 0.09 | 1.01 | 0.12 | 1.28 | 0.06 | 1.00 | 0.15 |
| RQVQPLAE | 6 | 69 | 76 | 0.69 | 0.11 | 0.64 | 0.18 | 0.71 | 0.11 | 0.63 | 0.14 | 0.85 | 0.10 | 1.07 | 0.05 | 0.88 | 0.09 |
| VRVPFTL | 5 | 77 | 83 | 0.12 | 0.04 | 0.13 | 0.04 | 0.06 | 0.08 | 0.09 | 0.08 | 0.15 | 0.05 | 0.24 | 0.03 | NaN | NaN |
| VRVPFTLD | 6 | 77 | 84 | 0.26 | 0.03 | 0.29 | 0.04 | 0.27 | 0.06 | 0.33 | 0.04 | 0.40 | 0.08 | 0.64 | 0.03 | 0.59 | 0.10 |
| LDSVARGATNPDLDD | 13 | 88 | 102 | 1.50 | 0.14 | 1.21 | 0.28 | 1.36 | 0.20 | 1.79 | 0.06 | 2.04 | 0.38 | 3.43 | 0.15 | NaN | NaN |
| DSVARGATNPDL | 10 | 89 | 100 | 1.25 | 0.04 | 1.12 | 0.12 | 1.25 | 0.07 | 1.39 | 0.10 | 1.69 | 0.29 | 3.12 | 0.17 | 2.27 | 0.23 |
| DSVARGATNPDLDD | 12 | 89 | 102 | 1.52 | 0.03 | 1.45 | 0.18 | 1.60 | 0.12 | 1.72 | 0.03 | 2.05 | 0.32 | 3.34 | 0.13 | 2.42 | 0.29 |
| AEDSAIFHG | 8 | 113 | 121 | 0.15 | 0.05 | 0.14 | 0.09 | 0.05 | 0.07 | 0.14 | 0.10 | 0.16 | 0.05 | 0.17 | 0.05 | -0.02 | 0.04 |
| HGWAQAGIKGIVDSTPHEAL | 18 | 120 | 139 | 2.27 | 0.08 | 2.15 | 0.51 | 2.54 | 0.23 | 2.84 | 0.10 | 3.25 | 0.41 | 6.04 | 0.09 | 5.14 | 0.58 |
| WAQAGIKGIVDSTPHEAL | 16 | 122 | 139 | 1.70 | 0.05 | 1.63 | 0.33 | 2.01 | 0.14 | 2.22 | 0.08 | 2.56 | 0.40 | 4.91 | 0.11 | 4.20 | 0.32 |
| IVDSTPHEAL | 8 | 130 | 139 | 0.68 | 0.07 | 0.72 | 0.12 | 0.86 | 0.08 | 0.97 | 0.04 | 1.07 | 0.19 | 2.39 | 0.12 | 1.85 | 0.21 |
| AVASVSDFPRAVL | 11 | 140 | 152 | 1.40 | 0.06 | 1.30 | 0.37 | 1.69 | 0.13 | 1.84 | 0.06 | 1.98 | 0.26 | 3.08 | 0.10 | 2.31 | 0.28 |
| SAADTLRKAGVTGPYA | 14 | 153 | 168 | 0.89 | 0.04 | 0.85 | 0.15 | 1.07 | 0.09 | 1.29 | 0.13 | 1.86 | 0.37 | 2.89 | 0.06 | 2.92 | 0.27 |
| SAADTLRKAGVTGPYAL | 15 | 153 | 169 | 0.85 | 0.07 | 0.82 | 0.14 | 1.08 | 0.09 | 1.25 | 0.08 | 1.75 | 0.44 | 2.85 | 0.08 | 2.92 | 0.28 |
| LVLGPKAYDDL | 9 | 169 | 179 | 0.47 | 0.06 | 0.44 | 0.11 | 0.59 | 0.08 | 0.62 | 0.05 | 0.75 | 0.10 | 1.27 | 0.06 | 1.04 | 0.15 |
| FAATQDGYPVAKQVQRL | 15 | 180 | 196 | 3.03 | 0.10 | 3.28 | 0.17 | 3.86 | 0.19 | 4.31 | 0.18 | 5.28 | 0.92 | 8.30 | 0.33 | 6.92 | 0.53 |
| VVDGPLVRA | 7 | 197 | 205 | 0.62 | 0.07 | 0.63 | 0.17 | 1.08 | 0.07 | 1.26 | 0.12 | 1.58 | 0.30 | 1.96 | 0.07 | 2.00 | 0.18 |
| VVDGPLVRANA | 9 | 197 | 207 | 1.08 | 0.06 | 1.20 | 0.13 | 1.70 | 0.09 | 2.02 | 0.08 | 2.59 | 0.18 | 3.26 | 0.10 | 2.84 | 0.19 |
| VVDGPLVRANAL | 10 | 197 | 208 | 1.09 | 0.04 | 1.02 | 0.28 | 1.83 | 0.18 | 2.01 | 0.19 | 2.75 | 0.21 | 3.84 | 0.10 | 3.53 | 0.31 |
| VVDGPLVRANALAGA | 13 | 197 | 211 | 1.58 | 0.06 | 1.45 | 0.44 | 2.50 | 0.27 | 2.86 | 0.09 | 3.76 | 0.32 | 5.20 | 0.15 | 5.17 | 0.29 |
| VVDGPLVRANALAGAL | 14 | 197 | 212 | 1.56 | 0.08 | 1.55 | 0.39 | 2.52 | 0.19 | 2.89 | 0.08 | 3.36 | 0.59 | 5.28 | 0.17 | 5.13 | 0.49 |
| NALAGAL | 6 | 206 | 212 | 0.46 | 0.05 | 0.50 | 0.14 | 0.70 | 0.09 | 0.77 | 0.06 | 1.10 | 0.18 | 1.92 | 0.07 | 2.14 | 0.07 |
| LSIGYAFHDRSKVEL | 14 | 229 | 243 | 0.28 | 0.13 | 0.31 | 0.13 | 0.38 | 0.15 | 0.34 | 0.16 | 0.93 | 0.23 | 1.85 | 0.12 | 1.59 | 0.19 |
| YAFHDRSKVEL | 10 | 233 | 243 | 0.48 | 0.07 | 0.41 | 0.11 | 0.39 | 0.11 | 0.42 | 0.20 | 0.75 | 0.12 | 1.18 | 0.29 | 0.95 | 0.30 |
| FVAESFT | 6 | 244 | 250 | 0.16 | 0.02 | 0.27 | 0.75 | 0.13 | 0.03 | 0.12 | 0.04 | 0.18 | 0.03 | 0.12 | 0.02 | NaN | NaN |
| FRVLEPGAA | 7 | 251 | 259 | 0.32 | 0.06 | 0.33 | 0.06 | 0.44 | 0.07 | 0.57 | 0.08 | 0.88 | 0.28 | 1.45 | 0.05 | 1.09 | 0.11 |

**Table S7: Deuterium uptake for each peptide and timepoint observed in HDX-MS of Empty-Enc.**

The deuterium uptake (in Da) is shown for each timepoint (10 s, 30 s, 2 mins, 5 mins, 4 hours and 24 hours) for each peptic peptide from Empty-Enc observed in HDX-MS. HDX-MS experiments were performed in triplicate and the standard deviation (SD) is shown for each time point. The number of exchangeable backbone amide hydrogens in each peptide is also shown (exchangers).

|  |  | **Empty-Enc deuterium uptake (Da)** | | | | | | | | | | | | | | | |
| --- | --- | --- | --- | --- | --- | --- | --- | --- | --- | --- | --- | --- | --- | --- | --- | --- | --- |
| **Sequence** | **Exchangers** | **Start** | **End** | **10s** | **10s SD** | **30s** | **30s SD** | **2 mins** | **2 mins SD** | **5 mins** | **5 mins SD** | **30 mins** | **30 mins SD** | **4 hrs** | **4 hrs SD** | **24 hrs** | **24 hrs SD** |
| LKRHLAPIVPDAW | 10 | 3 | 15 | 0.34 | 0.05 | 0.35 | 0.02 | 0.38 | 0.02 | 0.40 | 0.03 | 0.40 | 0.07 | 1.36 | 0.23 | 1.48 | 0.03 |
| LKRHLAPIVPDAWSA | 12 | 3 | 17 | 0.40 | 0.05 | 0.42 | 0.04 | 0.47 | 0.05 | 0.55 | 0.10 | 0.56 | 0.15 | 2.25 | 0.33 | 2.57 | 0.06 |
| EAKEIFQGHLAGRKLVD | 16 | 21 | 37 | 0.71 | 0.08 | 0.75 | 0.05 | 0.85 | 0.02 | 0.98 | 0.03 | 1.10 | 0.18 | 3.53 | 0.24 | 3.31 | 0.06 |
| EAKEIFQGHLAGRKLVDF | 17 | 21 | 38 | 0.65 | 0.05 | 0.68 | 0.04 | 0.84 | 0.01 | 1.07 | 0.04 | 1.21 | 0.25 | 3.58 | 0.20 | 3.29 | 0.06 |
| IFQGHLAGRKLVD | 12 | 25 | 37 | 0.37 | 0.04 | 0.40 | 0.03 | 0.43 | 0.03 | 0.47 | 0.09 | 0.55 | 0.09 | 1.81 | 0.28 | 1.60 | 0.06 |
| IFQGHLAGRKLVDF | 13 | 25 | 38 | 0.46 | 0.04 | 0.49 | 0.04 | 0.60 | 0.03 | 0.77 | 0.07 | 0.93 | 0.19 | 2.20 | 0.19 | 2.05 | 0.06 |
| FRGPFGWEY | 7 | 38 | 46 | 0.92 | 0.08 | 0.99 | 0.03 | 1.14 | 0.03 | 1.23 | 0.08 | 1.61 | 0.07 | 2.90 | 0.17 | 3.01 | 0.03 |
| YAAVNTGELRPIDDTPED | 15 | 46 | 63 | 1.96 | 0.15 | 2.15 | 0.11 | 2.63 | 0.08 | 2.93 | 0.10 | 3.45 | 0.51 | 5.76 | 0.20 | 5.71 | 0.17 |
| YAAVNTGELRPIDDTPEDVD | 17 | 46 | 65 | 2.07 | 0.12 | 2.24 | 0.10 | 2.76 | 0.10 | 3.12 | 0.12 | 3.51 | 0.53 | 6.70 | 0.22 | 6.61 | 0.15 |
| AAVNTGELRPIDDTPED | 14 | 47 | 63 | 1.90 | 0.12 | 2.14 | 0.07 | 2.60 | 0.04 | 2.94 | 0.10 | 3.25 | 0.44 | 5.25 | 0.18 | 5.18 | 0.17 |
| LRPIDDTPED | 7 | 54 | 63 | 1.07 | 0.05 | 1.22 | 0.01 | 1.41 | 0.04 | 1.54 | 0.09 | 1.65 | 0.25 | 2.42 | 0.14 | 2.36 | 0.03 |
| LRPIDDTPEDVD | 9 | 54 | 65 | 1.14 | 0.07 | 1.23 | 0.06 | 1.44 | 0.02 | 1.69 | 0.09 | 1.79 | 0.40 | 3.25 | 0.07 | 3.24 | 0.06 |
| MKLRQVQPLAE | 9 | 66 | 76 | 0.82 | 0.11 | 0.86 | 0.05 | 0.90 | 0.05 | 0.77 | 0.12 | 0.76 | 0.09 | 1.62 | 0.16 | 1.56 | 0.08 |
| KLRQVQPLAE | 8 | 67 | 76 | 0.79 | 0.08 | 0.84 | 0.06 | 0.88 | 0.07 | 0.85 | 0.08 | 0.88 | 0.10 | 1.60 | 0.20 | 1.48 | 0.08 |
| RQVQPLAE | 6 | 69 | 76 | 0.64 | 0.08 | 0.69 | 0.05 | 0.75 | 0.06 | 0.71 | 0.06 | 0.70 | 0.08 | 1.20 | 0.13 | 1.12 | 0.06 |
| VRVPFTL | 5 | 77 | 83 | 0.10 | 0.03 | 0.11 | 0.03 | 0.11 | 0.04 | 0.11 | 0.06 | 0.09 | 0.07 | 0.63 | 0.12 | 0.70 | 0.03 |
| VRVPFTLD | 6 | 77 | 84 | 0.24 | 0.03 | 0.24 | 0.02 | 0.30 | 0.02 | 0.35 | 0.04 | 0.42 | 0.09 | 1.01 | 0.08 | 1.04 | 0.03 |
| LDSVARGATNPDLDD | 13 | 88 | 102 | 1.37 | 0.16 | 1.48 | 0.11 | 1.64 | 0.08 | 1.55 | 0.21 | 1.61 | 0.49 | 3.68 | 0.29 | 3.53 | 0.14 |
| DSVARGATNPDL | 10 | 89 | 100 | 1.27 | 0.07 | 1.33 | 0.03 | 1.52 | 0.04 | 1.54 | 0.06 | 1.61 | 0.33 | 3.24 | 0.08 | 3.15 | 0.08 |
| DSVARGATNPDLDD | 12 | 89 | 102 | 1.48 | 0.12 | 1.59 | 0.04 | 1.80 | 0.02 | 1.95 | 0.08 | 2.13 | 0.27 | 3.82 | 0.16 | 3.61 | 0.12 |
| AEDSAIFHG | 8 | 113 | 121 | 0.12 | 0.04 | 0.13 | 0.05 | 0.12 | 0.04 | 0.11 | 0.08 | 0.09 | 0.08 | 0.18 | 0.11 | 0.13 | 0.03 |
| HGWAQAGIKGIVDSTPHEAL | 18 | 120 | 139 | 2.36 | 0.19 | 2.51 | 0.07 | 2.86 | 0.07 | 3.07 | 0.13 | 3.39 | 0.40 | 6.66 | 0.14 | 6.57 | 0.19 |
| WAQAGIKGIVDSTPHEAL | 16 | 122 | 139 | 1.76 | 0.18 | 1.89 | 0.14 | 2.19 | 0.14 | 2.44 | 0.15 | 2.74 | 0.42 | 5.65 | 0.19 | 5.45 | 0.19 |
| IVDSTPHEAL | 8 | 130 | 139 | 0.72 | 0.08 | 0.80 | 0.08 | 0.93 | 0.09 | 1.02 | 0.04 | 1.08 | 0.12 | 2.65 | 0.17 | 2.59 | 0.07 |
| AVASVSDFPRAVL | 11 | 140 | 152 | 1.52 | 0.08 | 1.66 | 0.04 | 1.94 | 0.05 | 2.09 | 0.04 | 2.19 | 0.19 | 3.42 | 0.11 | 3.38 | 0.10 |
| SAADTLRKAGVTGPYA | 14 | 153 | 168 | 0.93 | 0.07 | 1.05 | 0.05 | 1.32 | 0.04 | 1.53 | 0.06 | 1.71 | 0.31 | 3.39 | 0.10 | 3.37 | 0.06 |
| SAADTLRKAGVTGPYAL | 15 | 153 | 169 | 0.89 | 0.07 | 1.00 | 0.05 | 1.26 | 0.03 | 1.51 | 0.05 | 1.72 | 0.28 | 3.48 | 0.11 | 3.43 | 0.06 |
| LVLGPKAYDDL | 9 | 169 | 179 | 0.58 | 0.05 | 0.60 | 0.04 | 0.69 | 0.03 | 0.80 | 0.05 | 0.88 | 0.18 | 1.97 | 0.09 | 1.98 | 0.05 |
| FAATQDGYPVAKQVQRL | 15 | 180 | 196 | 3.62 | 0.16 | 3.96 | 0.19 | 4.97 | 0.10 | 5.40 | 0.04 | 6.34 | 0.09 | 8.32 | 0.12 | 8.39 | 0.10 |
| VVDGPLVRA | 7 | 197 | 205 | 0.80 | 0.06 | 0.93 | 0.06 | 1.39 | 0.06 | 1.68 | 0.04 | 1.83 | 0.21 | 2.58 | 0.15 | 2.70 | 0.06 |
| VVDGPLVRANA | 9 | 197 | 207 | 1.27 | 0.07 | 1.53 | 0.08 | 2.15 | 0.08 | 2.44 | 0.04 | 2.85 | 0.07 | 3.86 | 0.10 | 3.88 | 0.07 |
| VVDGPLVRANAL | 10 | 197 | 208 | 1.28 | 0.09 | 1.51 | 0.09 | 2.16 | 0.10 | 2.62 | 0.07 | 3.22 | 0.06 | 4.59 | 0.11 | 4.55 | 0.12 |
| VVDGPLVRANALAGA | 13 | 197 | 211 | 2.03 | 0.09 | 2.37 | 0.08 | 3.15 | 0.09 | 3.59 | 0.20 | 4.06 | 0.52 | 6.15 | 0.20 | 6.24 | 0.13 |
| VVDGPLVRANALAGAL | 14 | 197 | 212 | 2.08 | 0.10 | 2.41 | 0.10 | 3.24 | 0.06 | 3.58 | 0.10 | 4.26 | 0.10 | 6.39 | 0.18 | 6.41 | 0.16 |
| NALAGAL | 6 | 206 | 212 | 0.72 | 0.04 | 0.81 | 0.04 | 0.93 | 0.04 | 1.02 | 0.04 | 1.26 | 0.25 | 2.28 | 0.11 | 2.34 | 0.09 |
| LSIGYAFHDRSKVEL | 14 | 229 | 243 | 0.53 | 0.05 | 0.64 | 0.06 | 0.86 | 0.07 | 0.87 | 0.10 | 0.89 | 0.14 | 1.90 | 0.23 | 1.55 | 0.08 |
| YAFHDRSKVEL | 10 | 233 | 243 | 0.44 | 0.05 | 0.51 | 0.05 | 0.56 | 0.05 | 0.62 | 0.06 | 0.68 | 0.11 | 1.16 | 0.26 | 0.93 | 0.07 |
| FVAESFT | 6 | 244 | 250 | 0.09 | 0.01 | 0.09 | 0.01 | 0.10 | 0.01 | 0.11 | 0.03 | 0.07 | 0.02 | 0.12 | 0.03 | 0.10 | 0.01 |
| FRVLEPGAA | 7 | 251 | 259 | 0.29 | 0.03 | 0.38 | 0.55 | 0.50 | 0.03 | 0.72 | 0.04 | 0.89 | 0.20 | 1.46 | 0.10 | 1.36 | 0.03 |

### Table S8: gBlocks used in this study

| Construct | Sequence |
| --- | --- |
| Empty-Enc | GGCGAAGACAT**AATG**GATCTGCTGAAACGTCATCTGGCACCGATTGTTCCGGATGCATGGTCAGCAATTGATGAAGAAGCCAAAGAAATTTTTCAGGGCCATCTGGCAGGTCGTAAACTGGTTGATTTTCGTGGTCCGTTTGGTTGGGAATATGCAGCAGTTAATACCGGTGAACTGCGTCCGATTGATGATACACCGGAAGATGTTGATATGAAACTGCGTCAGGTTCAGCCGCTGGCCGAAGTTCGTGTGCCGTTTACCCTGGATGTTACCGAACTGGATAGCGTTGCACGTGGTGCAACCAATCCGGATCTGGATGATGTTGCCCGTGCAGCAGAACGTATGGTTGAAGCAGAAGATAGCGCAATTTTTCATGGTTGGGCACAGGCAGGTATTAAAGGTATTGTTGATAGCACACCGCATGAAGCACTGGCAGTTGCAAGCGTTAGCGATTTTCCGCGTGCAGTTCTGAGCGCAGCAGATACACTGCGTAAAGCCGGTGTTACCGGTCCGTATGCACTGGTTCTGGGTCCGAAAGCCTATGATGACCTGTTTGCAGCAACCCAGGATGGTTATCCGGTTGCAAAACAGGTGCAGCGTCTGGTTGTTGATGGTCCGCTGGTTCGTGCAAATGCCCTGGCAGGCGCACTGGTTATGAGCATGCGTGGTGGTGATTATGAACTGACCGTTGGTCAGGATCTGAGCATTGGTTATGCATTTCATGATCGTAGCAAAGTGGAACTGTTTGTGGCAGAAAGTTTTACCTTTCGTGTTCTGGAACCGGGTGCAGCCGTTCATCTGCGTTATGCATAA**AGGT**ATGTCTTCGTC |
| Loaded-Enc | Encapsulin: as above |
|  | EncFtn:  GGCGAAGACAT**AATG**AGCAGCGAACAGCTGCATGAACCGGCAGAACTGCTGAGCGAAGAAACCAAAAACATGCATCGTGCACTGGTTACCCTGATTGAAGAACTGGAAGCAGTTGATTGGTATCAGCAGCGTGCAGATGCCTGTAGCGAACCGGGTCTGCATGATGTTCTGATTCATAACAAAAACGAAGAGGTGGAACATGCAATGATGACCCTGGAATGGATTCGTCGTCGTAGTCCGGTTTTTGATGCACACATGCGTACCTACCTGTTTACCGAACGTCCGATTCTGGAATTAGAAGAAGAAGATACCGGTAGCAGCAGCAGCGTTGCGGCAAGCCCGACCAGCGCACCGAGTCATGGTAGCTTAGGTATTGGTAGCCTGCGTCAAGAAGGTAAAGAAGATTAA**AGGT**ATGTCTTCCGG |

BbsI restriction enzyme recognition sites are underlined, and overhang sequences are shown in bold.

### Table S9: Protein constructs used in this study.

| **Protein Construct** | **Protein** | **Sequence** | **Amino acids** | **Average Molecular weight (Da)** | **pI** | **Extinction coefficient (M^-1^ cm^-1^)** |
| --- | --- | --- | --- | --- | --- | --- |
| Empty-Enc | Encapsulin shell protein from  *H. ochraceum* | MDLLKRHLAPIVPDAWSAIDEEAKEIFQGHLAGRKLVDFRGPFGWEYAAVNTGELRPIDDTPEDVDMKLRQVQPLAEVRVPFTLDVTELDSVARGATNPDLDDVARAAERMVEAEDSAIFHGWAQAGIKGIVDSTPHEALAVASVSDFPRAVLSAADTLRKAGVTGPYALVLGPKAYDDLFAATQDGYPVAKQVQRLVVDGPLVRANALAGALVMSMRGGDYELTVGQDLSIGYAFHDRSKVELFVAESFTFRVLEPGAAVHLRYA**R** | 267 | 28968.84 | 4.88 | 26930 |
| Loaded-Enc | Encapsulin shell protein from  *H. ochraceum* | MDLLKRHLAPIVPDAWSAIDEEAKEIFQGHLAGRKLVDFRGPFGWEYAAVNTGELRPIDDTPEDVDMKLRQVQPLAEVRVPFTLDVTELDSVARGATNPDLDDVARAAERMVEAEDSAIFHGWAQAGIKGIVDSTPHEALAVASVSDFPRAVLSAADTLRKAGVTGPYALVLGPKAYDDLFAATQDGYPVAKQVQRLVVDGPLVRANALAGALVMSMRGGDYELTVGQDLSIGYAFHDRSKVELFVAESFTFRVLEPGAAVHLRYA | 266 | 28812.66 | 4.82 | 26930 |
|  | Encapsulated ferritin from  *H. ochraceum* | MSSEQLHEPAELLSEETKNMHRALVTLIEELEAVDWYQQRADACSEPGLHDVLIHNKNEEVEHAMMTLEWIRRRSPVFDAHMRTYLFTERPILELEEEDTGSSSSVAASPTSAPSHGSLGIGSLRQEGKED | 131 | 14799.39 | 4.70 | 13980 |

The additional Arg residue (bold, underlined) in the Hoch-Enc construct is a cloning artefact from the MoClo kit used.

### Table S10: Autoinduction media components

| **Media Component** | **Concentration** |
| --- | --- |
| Tryptone | 1 % (w/v) |
| Yeast Extract | 0.5 % (w/v) |
| Glycerol | 0.5 % (v/v) |
| Glucose | 0.05 % (w/v) |
| α-D-lactose | 0.2 % (w/v) |
| (NH_4_)_2_SO_4_ | 25 mM |
| KH_2_PO_4_ | 50 mM |
| Na_2_PO_4­_.7H_2_O | 50 mM |
| Magnesium Sulfate | 2 mM |

**Supplementary Movies**

**Supplementary Movie 1: Views of 3DVA component 0.**

Views of the trajectory of 3dVA component 0 showing the EM density maps rotating around the Y-axis, static, and as a cutaway to show movements of the EncFtn cargoes.

**Supplementary Movie 2: Views of 3DVA component 1.**

Views of the trajectory of 3dVA component 0 showing the EM density maps rotating around the Y-axis, static, and as a cutaway to show movements of the EncFtn cargoes.

**Supplementary Movie 3: Views of 3DVA component 2.**

Views of the trajectory of 3dVA component 0 showing the EM density maps rotating around the Y-axis, static, and as a cutaway to show movements of the EncFtn cargoes.
